# The human and mouse synaptome architecture of excitatory synapses show conserved features

**DOI:** 10.1101/2020.01.17.910547

**Authors:** Olimpia E. Curran, Zhen Qiu, Colin Smith, Seth G. N. Grant

## Abstract

Large-scale mapping of the location of synapses and their molecular properties in the mouse has shown that diverse synapse types are spatially distributed across the brain. The diversity of synapses is known as the synaptome and the spatial distribution as the synaptome architecture. Synaptome maps in the mouse show each brain region has a characteristic compositional signature. The signature can store behavioral representations and is modified in mouse genetic models of human disease. The human synaptome remains unexplored and whether it has any conserved features with the mouse synaptome is unknown.

As a first step toward creating a human synaptome atlas, we have labelled and imaged synapses expressing the excitatory synapse protein PSD95 in twenty human brain regions in four phenotypically normal individuals. We quantified the number, size and intensity of approximately a billion individual synaptic puncta and compared their regional distributions. We found that each region showed a distinct signature of synaptic puncta parameters. Comparison of brain regions showed the synaptome of cortical and hippocampal structures were similar but distinct to the synaptome of cerebellum and brainstem. Comparison of human and mouse synaptome revealed conservation of synaptic puncta parameters, hierarchical organization of brain regions and network architecture. These data show that the synaptome of humans and mouse share conserved features despite the 1000-fold difference in brain size and 90 million years since a common ancestor. This first draft human synaptome atlas illustrates the feasibility of generating a systematic atlas of the human synaptome in health and disease.

## Introduction

For over a century synapses have been recognised as a hallmark of brain anatomy, yet there are no reference resources that report the number and distribution of synapse types at the single synapse resolution across human brain regions. The importance of understanding the molecular anatomy of synapses is underscored by the remarkable growth in knowledge about the importance of synaptic proteins in disease. In 2000 there were three human genetic disorders known that disrupted proteins in excitatory synapses (Husi *et al*., 2000) and by 2011 this had been extended to 133 common and rare brain disorders (Bayes *et al*., 2011). Schizophrenia, depression, autism spectrum disorders and intellectual disability are all caused by gene mutations targeting synapse proteins (Husi *et al*., 2000; Fernandez *et al*., 2009; Bayes *et al*., 2011; Nithianantharajah *et al*., 2013; Fromer *et al*., 2014; Purcell *et al*., 2014; Fernandez *et al*., 2017; Howard *et al*., 2018; Satterstrom *et al*., 2019). The expression of synapse proteins is also altered in neurodegenerative disorders, brain injury (including stroke and traumatic brain injury), neuroinflammatory conditions (including multiple sclerosis and autoimmune encephalopathies) and addiction (including alcohol and cocaine). It is now accepted that synapse molecular pathology is a feature of most brain disorders.

Mammalian synapses contain thousands of evolutionarily conserved proteins (Bayés *et al*., 2011; Bayes *et al*., 2012; Bayés *et al*., 2017) and these are differentially distributed across the brain (Emes *et al*., 2008; Hawrylycz *et al*., 2012; Roy *et al*., 2018a; Roy *et al*., 2018b). A transformative concept, that has emerged from the capacity to image synaptic proteins in individual synapses, is that there is a far greater diversity than could be anticipated from historical studies of synapse morphology, neurotransmitters and physiology (O’Rourke *et al*., 2012; Zhu *et al*., 2018a). Driven by the ability to systematically characterise the molecular composition of individual synapses across the whole brain, the term ‘synaptome’ has been introduced to describe the catalogue of synapses in the brain or part thereof (Zhu *et al*., 2018a).

To map the mouse synaptome, we developed a suite of technologies into a pipeline called SYNMAP: endogenous synaptic proteins are labelled (with genetic tags or immunolabelling) and tissue sections are imaged with high-speed spinning disc confocal microscopy, then sophisticated image analysis tools exploiting computer vision and machine-learning detect individual synapses and measure their molecular and morphological properties, then classify the synapses and map their spatial locations in brain atlases. Mapping the spatial location of synapse types and subtypes in mice reveals there is synaptome architecture of the brain (Zhu *et al*., 2018a). Synapse diversity is observable at all scales – individual dendrites, cell types, brain regions to the global systems level (Zhu *et al*., 2018a). Studies of genetic disorders in mice show that synapse diversity is modified and the synaptome architecture, suggesting that this could be a common feature of synaptic diseases.

The synaptome architecture of human brains is poorly understood and as a first step toward creating a human synaptome atlas, we recognized the need to overcome several challenges. First, the need to develop robust molecular labelling of excitatory synapses in post-mortem brain tissue using immunohistochemistry (IHC) suitable for analysis using the SYNMAP pipeline. We chose to label excitatory synapses using antibodies to PSD95, which is an abundant postsynaptic protein known to play a key role in physiology, behaviour and disease (Migaud *et al*., 1998; Bayés *et al*., 2011). To optimize and validate our IHC labelling of PSD95, we took advantage of two lines of genetically modified mice that we had previously characterized: mice lacking PSD95 (PSD95^-/-^) (Migaud *et al*., 1998) and mice expressing PSD95 fused to green fluorescent protein (PSD95eGFP) (Zhu *et al*., 2018a). PSD95^-/-^ and PSD95eGFP mice serve as negative and positive controls, respectively, for antibody staining. Second, given that the human brain is 1000-fold larger than that of the mouse, we needed to explore the feasibility of acquiring single-synapse resolution data across the major brain regions. Although PSD95 immunostaining has been reported in specific human brain regions, including frontal cortex, caudate nucleus, putamen and hippocampus (Glantz *et al*., 2007; Fourie *et al*., 2014; Morigaki & Goto, 2015), there is no single comprehensive study of PSD95 distribution across a number of human brain areas at single-synapse resolution using a high-throughput quantitative analysis.

Here we report a detailed qualitative and quantitative survey of the distribution and diversity of PSD95-labelled synapses throughout 20 brain areas from four phenotypically normal individuals. We describe the synaptome architecture of human brain regions and analyze regional similarity within the human brain and between human and mouse brain. We discuss the utility of this dataset for studies of the normal and diseased brain and the logistical considerations for large-scale human synaptome mapping.

## Materials and methods

### Consent

Post-mortem human brain tissue was obtained from the Medical Research Council UK (MRC) funded University of Edinburgh Brain Bank (EBB). All procedures involving post-mortem human brain tissue were approved by East of Scotland Research Ethics Service (16/ES/0084). Informed written consent was obtained in releation to each subject.

### Human subjects

Human brains from four control subjects (three males, one female; mean age 51 ± 6.9 (standard deviation) years) were used to image PSD95 using laser scanning confocal microscopy (LSCM). Details of the human subjects are listed in Table 1. None of the control subjects had a history of dementia, neurological or psychiatric disorders at the time of death. Human brain tissue donated to the EBB was processed according to departmental protocols. Briefly, formalin-fixed paraffin-embedded (FFPE) 4 μm tissue sections were stained with hematoxylin and eosin (H&E) for routine neuroanatomical and neuropathological surveys. Selected sections were stained with a panel of antibodies, which included phosphorylated tau for neuronal and glial inclusions, beta-amyloid for vascular and parenchymal amyloid deposition, ubiquitin for ubiquitinated aggregates and/or alpha-synuclein for Lewy body pathology. Gross anatomical and microscopic examinations of the brain tissue revealed only low age-related neuropathological changes in these control brains (Table S1).

**TABLE 1.**
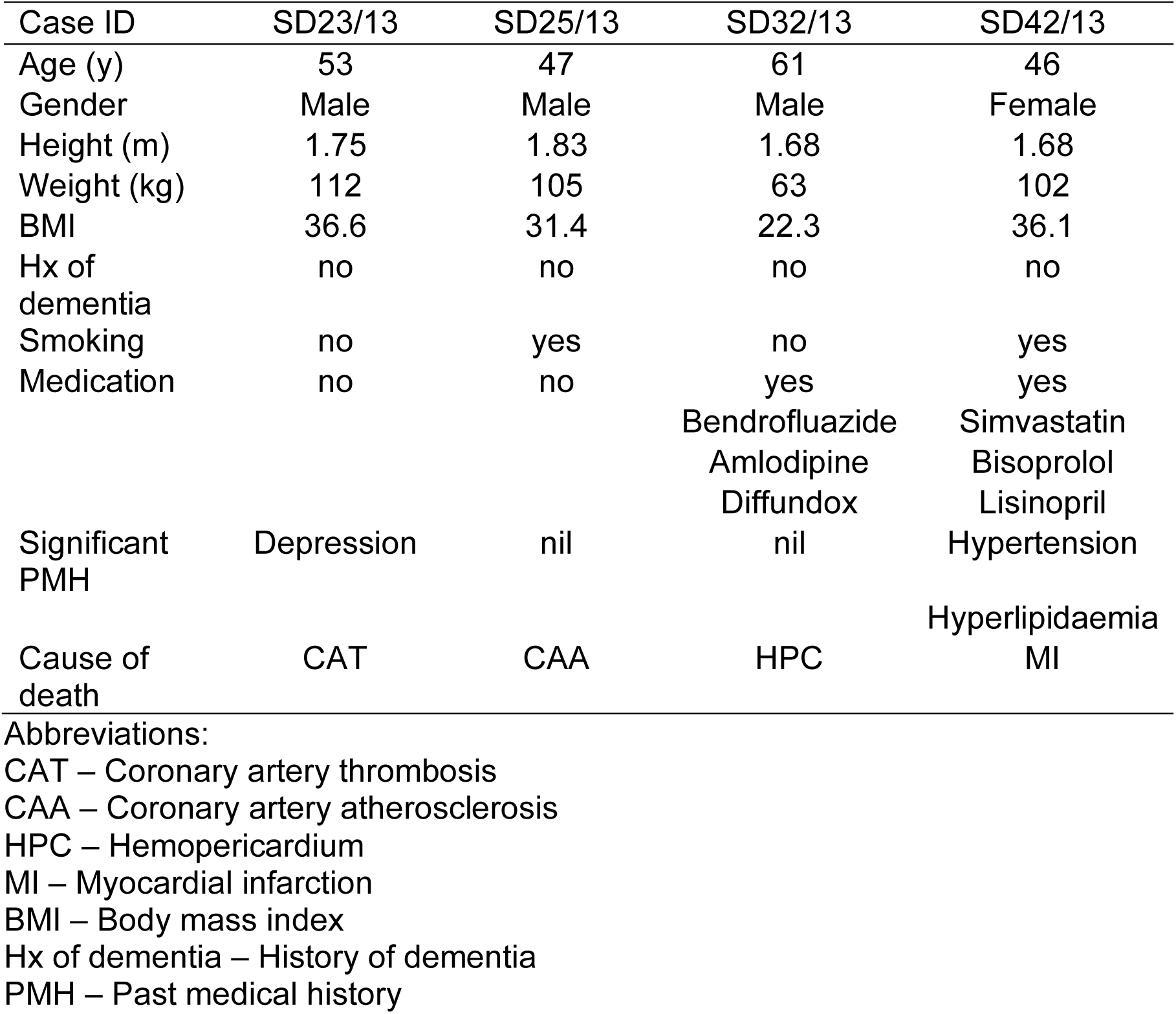
Details of the human control subjects used in this study.

### Post-mortem procedures

At post-mortem examination brains were macroscopically dissected into 1 cm-thick coronal slices and sampled according to the EBB protocol (Samarasekera *et al*., 2013). Small tissue blocks were immediately fixed in 10% formaldehyde for 24-72 h prior to further tissue processing and embedding in paraffin wax. The blocks were sampled from 13 neocortical areas (six frontal, four temporal, one parietal, two occipital), one allocortical area (hippocampus), two subcortical grey matter areas (thalamus and caudate nucleus) and four infratentorial regions (midbrain, pons, medulla and cerebellum) (Table 2). The 13 neocortical areas are defined as Brodmann areas, and include sensory visual regions (BA17 and BA19), sensory auditory region (BA41/42), motor region (BA4), premotor region (BA6/8), associative frontal regions (dorsolateral BA9 and BA46, ventrolateral BA44/45, and orbital BA11/12), associative temporal regions (BA20/21 and BA38), associative parietal region (BA39) and associative occipitotemporal region (BA37). The pathological changes in neurodegenerative diseases are present throughout the brain parenchyma; however, careful selection of representative areas allows efficient screening of tissue for any potential disease entities (King *et al*., 2013). The choice of the anatomical regions examined was additionally guided by the availability of all 20 blocks for the four control cases. Tissue blocks were macroscopically dissected by experienced pathologists and the selection of blocks closely followed topographical brain anatomy using the patterns of sulci and gyri. Regions of interest were further identified using an atlas of human brain (Mai *et al*., 2015). Nissl-stained sections from blocks immediately adjacent (where possible) to those used for IHC were examined to assess and confirm the distinctive cytoarchitectural features of various areas. The absence of significant neuropathological findings was further confirmed by routine examination of immunohistochemical and special stains available from the Department of Academic Neuropathology, Edinburgh University.

**TABLE 2.**
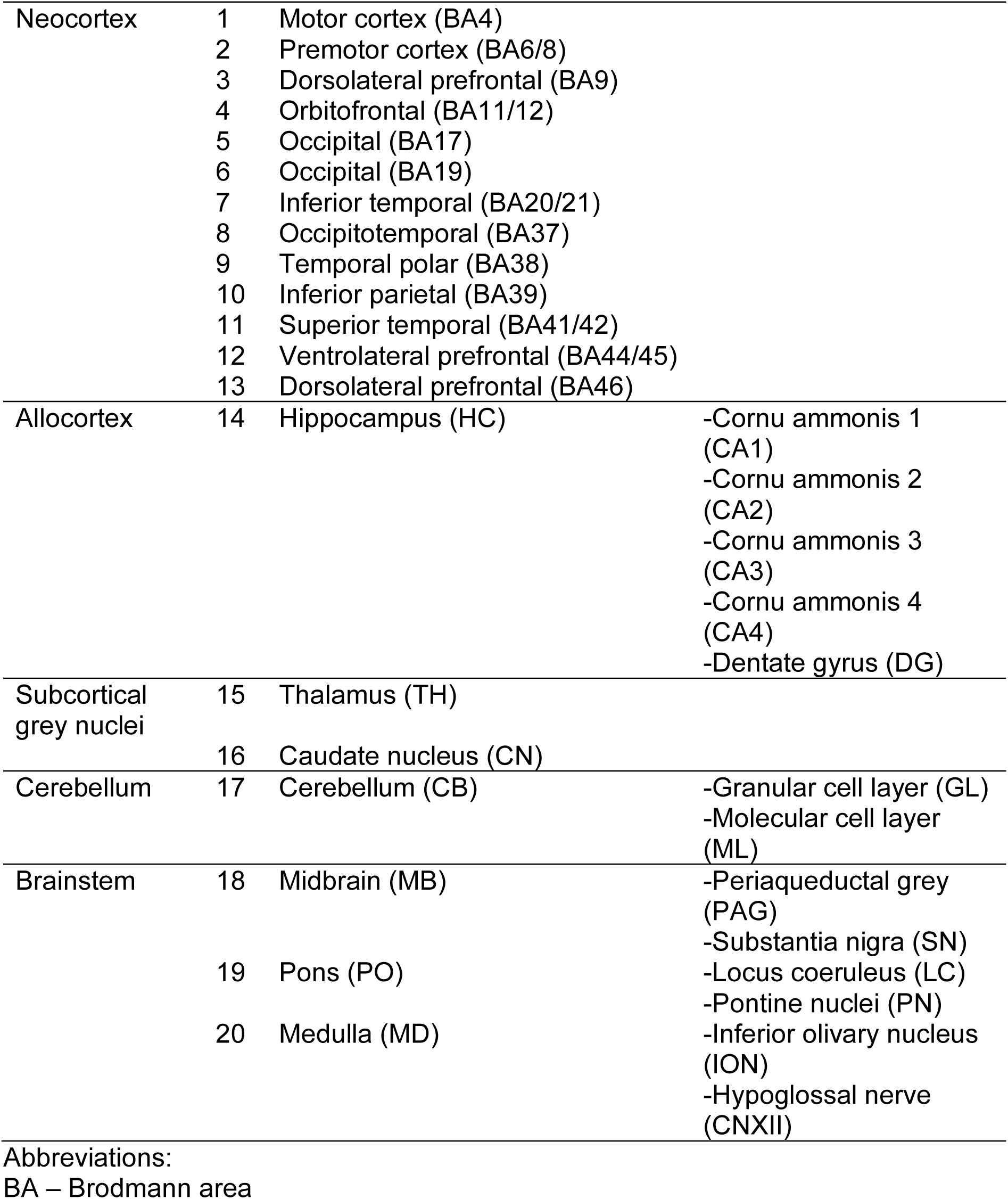
Human brain areas analysed in this study.

### Fluorescence immunohistochemistry

FFPE human brain sections were dewaxed and dehydrated as per the EBB neuropathology laboratory protocol. Briefly, sections were dewaxed in two 3-min xylene washes followed by four 3-min graded alcohol washes, comprising two 74°OP (IMS99%) alcohol washes and two 70% alcohol washes. This was followed by a 15-min wash in saturated alcoholic picric acid to remove any formalin sedimentation and a final wash in running tap water for 15 min. Antigen and heat retrieval techniques were used to enhance cell surface staining, antigen detection from antigen-masked FFPE sections and reduction of non-specific background staining. The slides were placed in 250 ml freshly made antigen retrieval solution (0.1 M sodium citrate buffer pH 6.0, Fisher Scientific) and pressure cooked (A. Menarini Diagnostics) at 125°C for 30 s. After cooling in tap water, sections were washed once with PBS and blocked with 5% bovine serum albumin (BSA, Sigma-Aldrich) in 1x Tris-buffered saline (TBS, Fisher BioReagents) containing 0.2% Triton X-100 (Sigma-Aldrich) for 1 h at room temperature (RT). Sections were then incubated overnight at 4°C with 16 ng/ml mouse anti-PSD95 monoclonal IgG2a (NeuroMab) primary antibody diluted (1:250 of 4 μg/ml stock) in TBS containing 3% BSA and 0.2% Triton X-100. After three 5-min washes with TBS and 0.2% Triton X-100, secondary antibodies Alexa Fluor 546 goat anti-mouse IgG2a (γ2a) (Molecular Probes) were added at 1:500 and sections incubated for 2 h at RT. Each IHC staining round was accompanied by nuclear cell counterstaining with 4’,6-diamidino-2-phenylindole (1μg/ml DAPI (Sigma) diluted 1:1000 in PBS). Finally, the slides were washed three times with TBS and 0.2% Triton X-100. For elimination or reduction of lipofuscin-like autofluorescence, sections were subsequently treated with Autofluorescence Eliminator Reagent (AER, Millipore Chemicon International). Lipofuscin pigment accumulates in the cytoplasm of brain cells and can hamper fluorescence microscopy owing to its broad excitation and emission spectra. AER is a commercially available Sudan Black (SB)- based reagent that has been successfully used in studies using human tissue (Blazquez-Llorca *et al*., 2010). SB has been reported to provide the best compromise between the reduction of lipofuscin-like fluorescence and maintenance of specific fluorescent labels (Schnell *et al*., 1999). Briefly, the sections were first immersed in 70% ethanol for 5 min, followed by the AER for 30 s. Slides were washed in 70% ethanol for a further 3 min before being mounted with mounting medium (Mowiol with 1,4-diazabicyclo[2.2.2]octane (DABCO), Sigma-Aldrich) and left to dry overnight.

### Validation of PSD95 IHC with PSD95eGFP mouse

Qualitative and quantitative comparisons with a selection of images from PSD95eGFP mice were made to validate the human PSD95 staining. At first, the representative IHC images of coronal sections from several mouse brain areas were acquired using the same LSCM and with the same parameter set-up. Then, a direct quantitative comparison of PSD95eGFP mouse versus human results was made for three wild-type (WT) male mice (aged 18 months) and three human male control subjects from ten brain regions. The PSD95eGFP mouse data were acquired with a spinning disk microscope (SDM) and analysed using the Ensemble analysis as previously described (Zhu *et al*., 2018a).

### Image acquisition

To resolve individual synaptic puncta, antibody-labelled brain tissue sections were captured using a Zeiss laser scanning confocal microscope (LSCM510) using a Zeiss Plan-Apochromat 63x oil-immersion objective lens (NA 1.4) with a frame size of 1024 x 1024 pixels, a pixel dimension of 46 x 46 nm and a depth of 8 bit. All raw images were obtained with x3.1 zoom as stacks of six image planes with *z*-step of 130 nm pixels and a total length of axial plane 780 nm pixels. The total area used for synaptic density determination varied depending on region, but for cortical areas 18 images randomly distributed per Brodmann area (three per cortical layer) per subject were acquired. A total of 1510 confocal images were acquired, of which 21 (1.4%) were excluded due to technical problems, including too low signal-to-noise ratio precluding adequate image analysis, a lack of alignment in *z*-stacks, and images with overexposed synaptic puncta. It is estimated that more than half a billion individual synaptic puncta were identified and counted in this study. Images and figure montages were prepared in Adobe Photoshop and Adobe Illustrator. Where necessary, images were adjusted for brightness for correct display, but no other corrections were made.

For the visualisation of the whole-mounted brain sections, a Zeiss Axio Scan.Z1 Slide Scanner (Carl Zeiss) with a 20x lens (NA 0.8) was used. Images were obtained using sections stained with secondary antibodies conjugated to Alexa Fluor 546, DAPI and Alexa Fluor 633. Pixel resolution was 0.325 x 0.325 μm and image brightness is in a depth of 16 bits. The ZEN lite 2012 software (Zeiss) was used with the scanner for image acquisition and adjustment. Coronal brain section images were background and contrast adjusted to provide consistent comparison of PSD95 expression. All control images were directly comparable with their negative controls using the same settings. The slide scanner-captured images were used only for qualitative comparisons. The slide scanner was also used to capture Nissl-stained material for direct assessment of tissue integrity and cytoarchitecture of sections used in this study.

### Image analysis

Image analysis of the human synapse images includes image puncta detection, segmentation and parameter quantifications. The detection and segmentation of individual puncta were done by ensemble learning, a machine learning approach developed in house (Zhu *et al*., 2018a). Ensemble learning of image detection requires an annotated image set from the LSCM dataset for algorithm training. The raw tile images were firstly selected from the whole data set using random bootstrapping. PSD95-labelled individual synaptic puncta in each of the images were then manually located using the Fiji plugin ‘Cell Counter’ (Schindelin *et al*., 2012) by two independent experts. Half of the annotated data were used for training and the other half for testing of thee trained detector. A K folder strategy was also used for validation of the training.

The second step in the ensemble image analysis involves algorithm training of synaptic puncta detector, which is based on a multi-resolution image feature detection and supervised machine-learning technique. A multi-resolution and multi-orientation version of 2^nd^-order nonlocal derivative (NLD) to calculate intensity differences, or image features, was developed for each of all individual puncta. For instance, for an image of PSD95 IHC, a group of 109 feature images were calculated per punctum. These intensity differences were assembled as feature vectors of each individual punctum. Details of the algorithm training can be referred to our previous work (Zhu *et al*., 2018a). Once training was finished, the trained ensemble detector was applied to automatically detect puncta in the LSCM images. Once being detected, the object is then segmented by thresholding the pixel intensity of the puncta. The threshold was calculated adaptively for each punctum as the 18% of the maximal intensity of the punctum after subtracting the background intensity.

Different puncta parameters were measured after the detection and segmentation. In particular, the mean punctum intensity is a measurement of the relative amount of PSD95 protein within the PSD, since the absolute number of PSD95 molecules cannot be resolved using confocal imaging. Nevertheless, the mean punctum intensity provides a good insight into the differences in average packing of PSD95 between areas or individuals. Besides, the puncta size is a measurement of the relative area of PSD95-positive synapses. It is calculated as the number of pixels within the segmented puncta and converted in μm^2^. The final measurement provided by ensemble image analysis is the puncta density, i.e. the number of detected puncta per 100 μm^2^.

### Statistical analysis

Three PSD95 punctum parameters, synaptic puncta density (100 μm^2^), puncta intensity (A.U.) and size (μm^2^), were used for the statistical study. The punctum size is only an approximation when the square root of the results was taken, as majority of puncta are not perfectly spherical. The parameters were plotted with the Tukey-style as box and whisker plots (Krzywinski & Altman, 2014). Normality of LSCM and SDM data distribution was analyzed using D’Agostino-Pearson omnibus normality test. For a heatmap generation, the data were reorganized using an agglomerative hierarchical clustering algorithm and standardized using the z-score normalization. A similarity matrix in Figure 13 was calculated to quantify the synaptome network architecture in the human brain. Each row/column in the matrix represents a brain region/subregion. The rows/columns are resorted to indicate the brain anatomy: anatomically close regions are placed in the adjacent rows/column in the matrix. Elements of the matrix are the similarity and calculated using the Euclidean distances of the three synaptic parameters between the two regions. It ranges from zero to one, the latter of which indicates identical synaptome of the two regions. Similarity matrix was plotted as a heatmap where the elements were colour coded.

## Results

The study is divided into four phases: i) the optimization of synaptic labelling with antibodies to PSD95, ii) the systematic acquisition of image data from 20 brain regions with a detailed description of the synaptic anatomy, iii) an analysis of the populations of synapses in the brain areas, and iv) a comparison of the synaptome architecture of human and mouse.

### Optimization of PSD95 immunostaining

To identify optimal IHC conditions we used four complementary approaches. First, we used the standard IHC controls including the presence and absence of primary and secondary antibodies on human tissue (Figure 1). Strong punctate staining was observed with PSD95 antibodies and abolished when either primary or secondary antibodies were absent. Second, to confirm that the PSD95^+^ puncta observed were synaptic, we labelled the excitatory presynaptic terminals with synapsin 1 and synaptophysin (Figure 2) (Koffie *et al*., 2009) and detected the characteristic juxtaposed labelling in 91.5% and 87.8% of synapses, respectively. Third, we performed PSD95 IHC on brain sections from *Psd95* knockout mice and confirmed the absence of positive signal (Figure 3). Fourth, we compared the puncta distribution of multiple human brain regions with the corresponding regions in mice expressing PSD95eGFP, which revealed similar synaptic puncta distributions in these areas (Figure 4). We screened eight commercially available antibodies and found best results with anti-PSD95 monoclonal IgG2a (NeuroMab) at 16 ng/ml (a 250-fold dilution of the 4 μg/ml stock solution).

**FIGURE 1.**
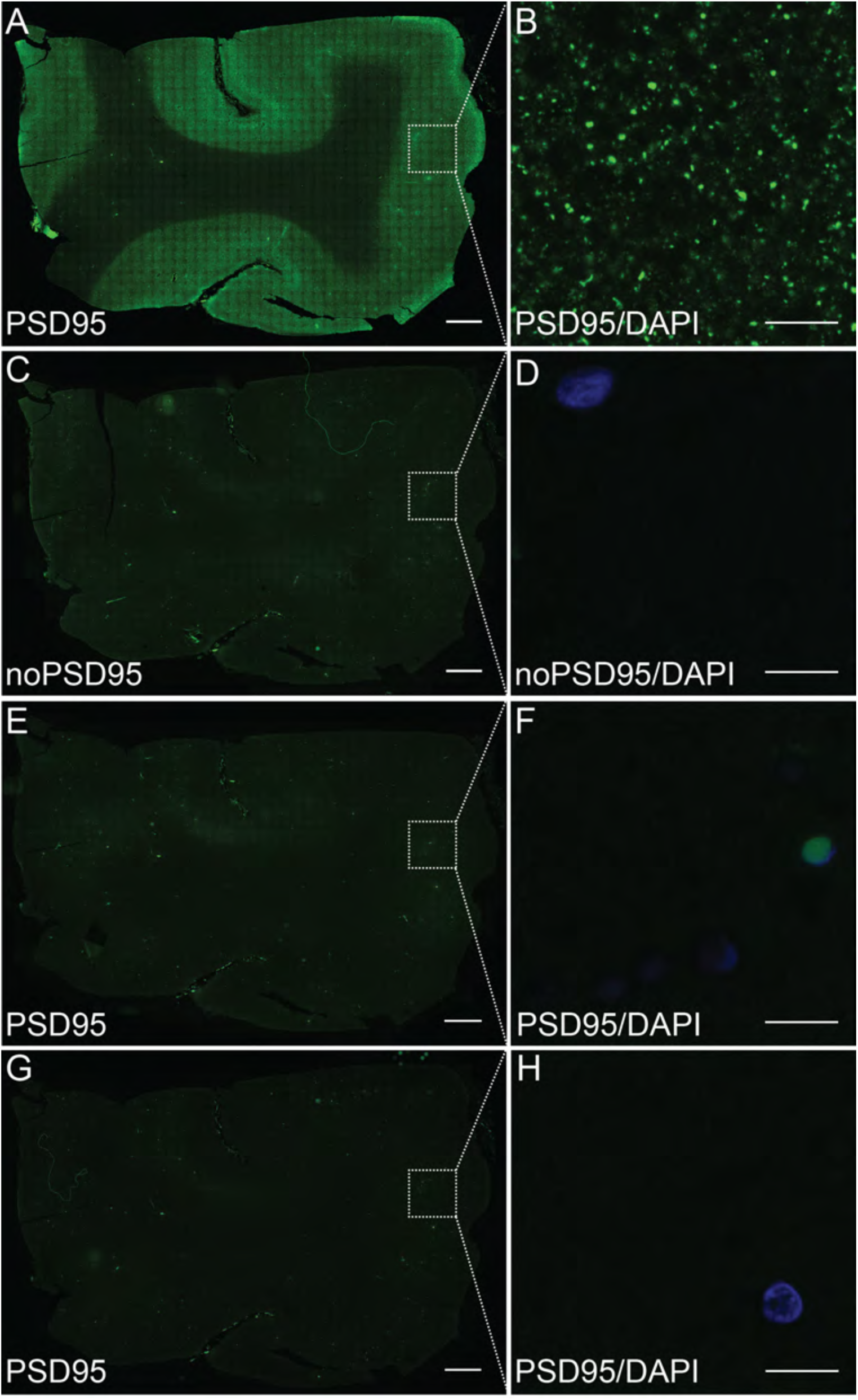
PSD95 antibody specificity in human post-mortem tissue. (A, B) Normal punctate PSD95 immunostaining. (C, D) Punctate PSD95 staining was abolished when the primary antibody was omitted. (E, F) No PSD95 punctate staining was detected when the secondary antibody was omitted. (G, H) No PSD95 punctate staining was detected when the secondary antibody came from another species. Scale bars: 1 mm in A, C, E, G; 10 μm in B, D, F, H.

**FIGURE 2.**
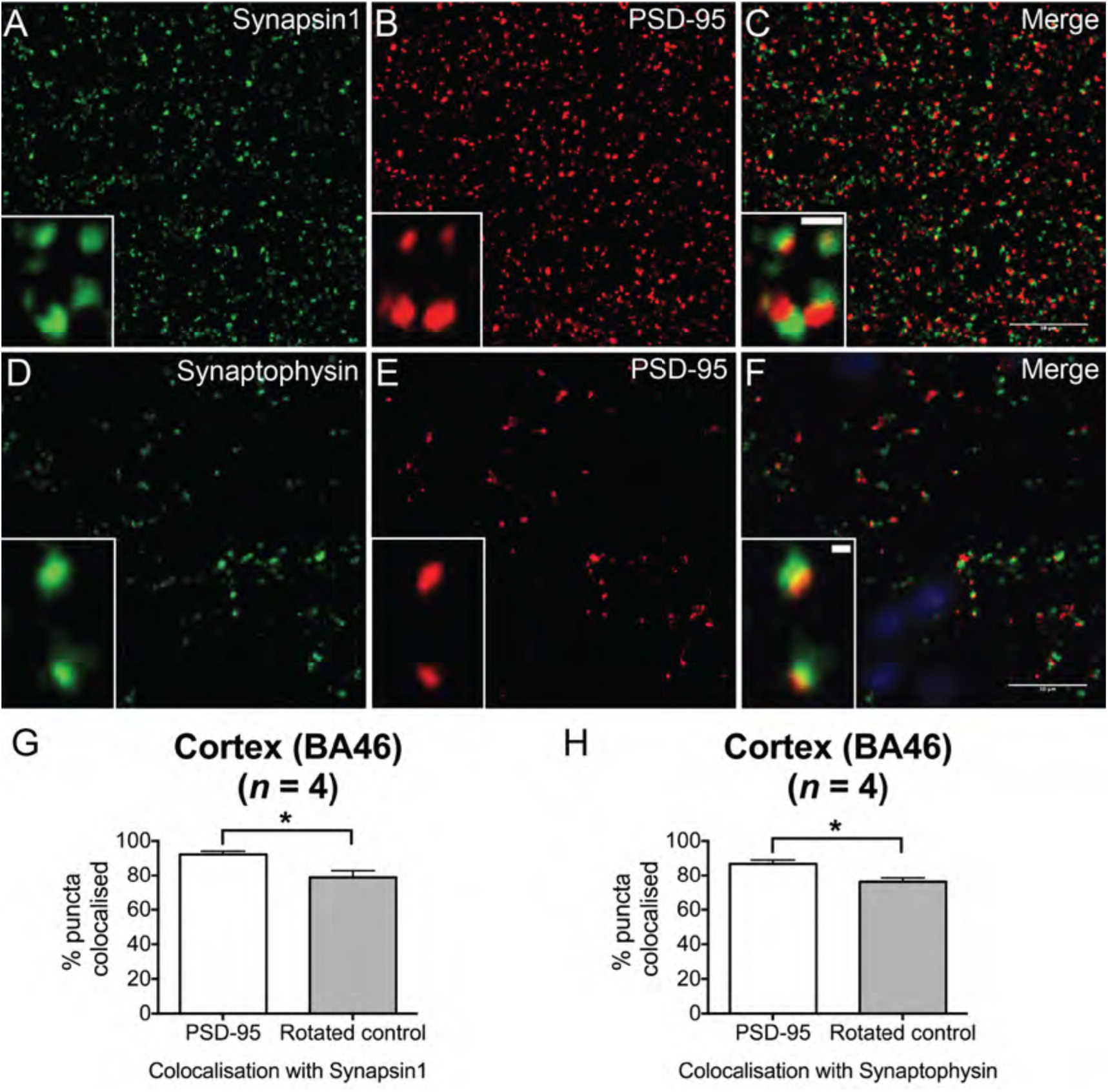
PSD95 antibody as a reliable excitatory postsynaptic marker. (A-C) PSD95 co-stained with synapsin 1 antibody. (D-F) PSD95 co-stained with synaptophysin antibody. Both, synapsin 1 and synaptophysin are examples of presynaptic markers and these images demonstrate that postsynaptic PSDs are opposed by presynaptic terminals. Insets (C, F) demonstrate the typical immunofluorescent ‘snowman’ appearance of colocalised pre- and postsynaptic markers. Overall, the majority of PSD95 puncta are juxtaposed to presynaptic terminals. Scale bars: 10 μm in C, F; 1 μm in insets. (G, H) Quantification of PSD95-positive puncta colocalisation with synapsin 1 (G) and synaptophysin (H). The rotated control shows the degree of random colocalisation obtained by clockwise rotation of the synapsin 1 (G) or synaptophysin (H) images. The colocalisation with either presynaptic marker is significantly higher than expected by chance. Data are mean ± SD; *P ≤ 0.05, Mann-Whitney U-test.

**FIGURE 3.**
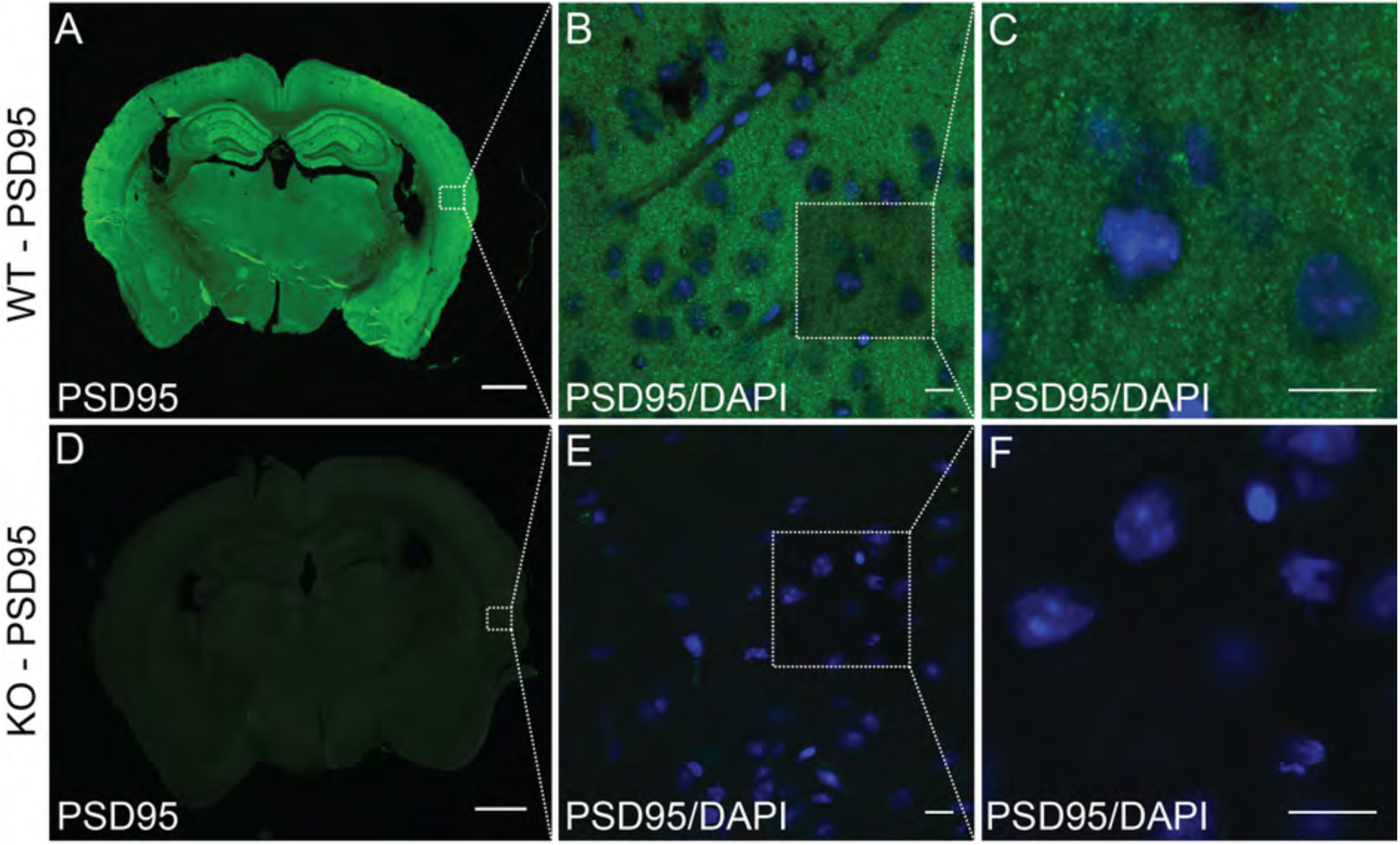
PSD95 antibody staining in wild-type and *Psd95* knockout mice. (A-C) PSD95 staining in a wild-type (WT) mouse. (D-F) PSD95 staining is abolished in *Psd95* knockout (KO) mouse. (A, D) low magnification (20x). (B, C, E, F) high magnification (63x). Scale bars: 1 mm in A, D; 10 μm in B, C, E, F.

**FIGURE 4.**
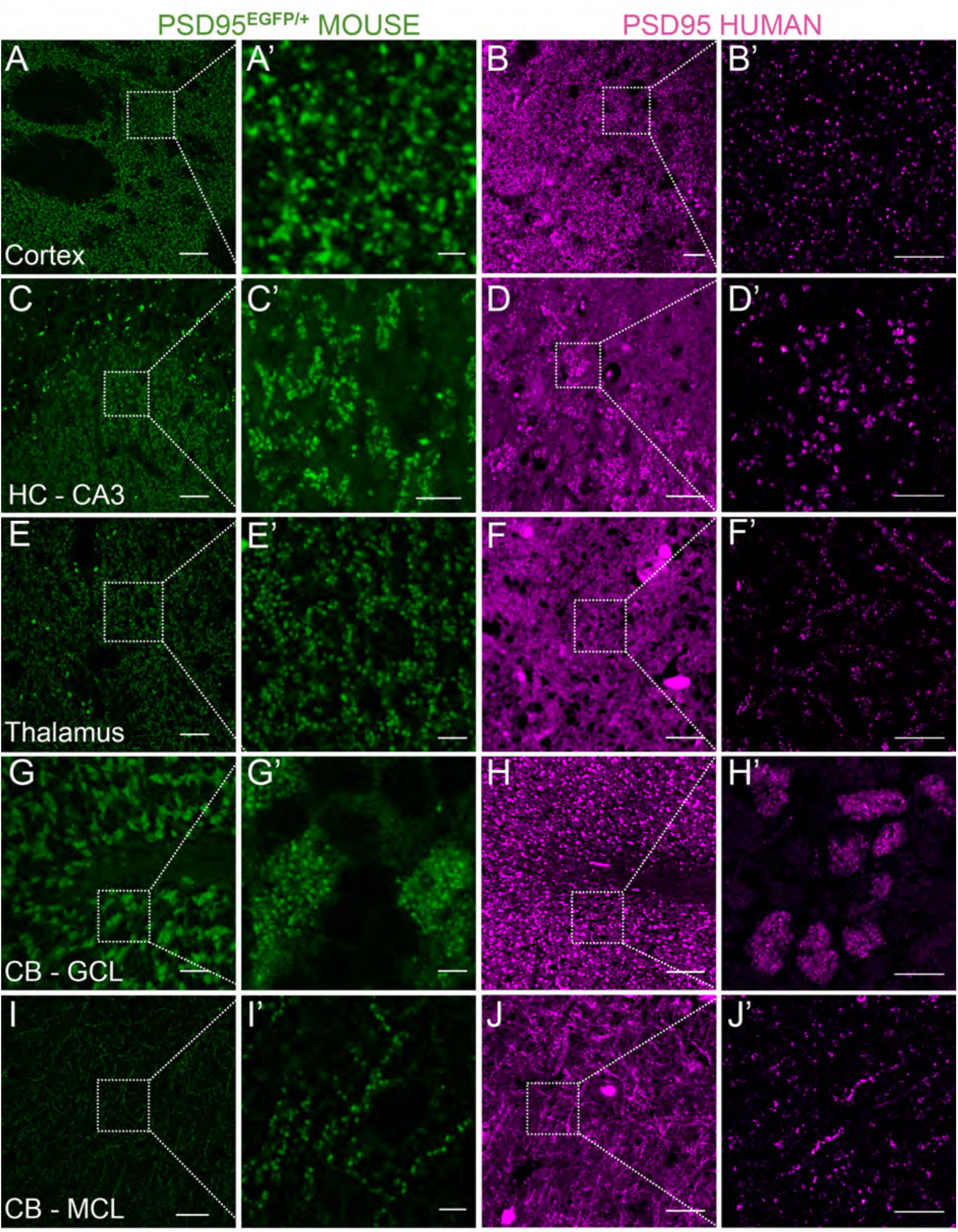
Comparison of PSD95 expression between human and mouse brain. Similarity of PSD95 immunofluorescence patterns between human and mouse. Panels in green: tissue from mutant knock-in mouse examined using direct fluorescence microscopy of PSD-EGFP fusion protein in the adult *Psd95*^EGFP/+^ heterozygote. Panels in magenta: human tissue stained with PSD95 antibody using indirect immunofluorescence. (A-B’) Similar patterns of staining were observed in human and mouse cortex. (C-D’) Similar patterns of staining were observed in hippocampal CA3 region. However, the staining was present in two different subregions: in mouse in stratum lucidum (C, C’), but in human in stratum pyramidale (D, D’). (E-F’) Similar patterns of staining were observed in thalamus. (G-H’) Similar patterns of staining were observed in cerebellum granular cell layer. (I-J’) Similar patterns of staining were observed in cerebellar molecular cell layer. HC, hippocampus; CA3, cornu ammonis; CB, cerebellum; GCL, granular cell layer; MCL, molecular cell layer. Scale bars: 20 μm in A, C, E, G, I; 2 μm in A’, E’, G’, I’; 5 μm in C’; 30 μm in B, D, J; 100 μm in F, H; 10 μm in B’, D’, F’, H’, J’.

### Human brain regions show different distributions of excitatory synapses

In order to describe the distribution patterns of the PSD95^+^ synapses in the human brain, we performed IHC on post-mortem material from a selection of 20 regions. Representative images from these selected brain areas are shown in Figure 5. PSD95 expression was detected as fluorescent puncta (< 1 μm in size) in all regions. In general, the PSD95 puncta appeared intense and densely packed in the neocortex and the hippocampus. However, even within the neocortex, different areas displayed unique combinations of bright, dim, large and small puncta. In the subcortical structures, the thalamus and caudate nucleus, the puncta tended to show moderate densities and intensities. A high density of intense PSD95 puncta was found in some parts of the cerebellum, such as the granular cell layer, but not others, such as the cerebellar white matter. Both the hippocampus and cerebellum showed distinct subregional distribution patterns of PSD95 puncta, as described below. Although present within the brainstem, the synaptic puncta density was low in the midbrain, pons and medulla. Synaptic puncta were almost absent from the cortical and the cerebellar white matter (not shown). Thus, human brain regions show diversity in the densities, intensities and size of PSD95 puncta. In the following sections, we describe some of the diversity observed within the hippocampal formation, the deep grey nuclei (caudate and thalamus), cerebellum and brainstem.

**FIGURE 5.**
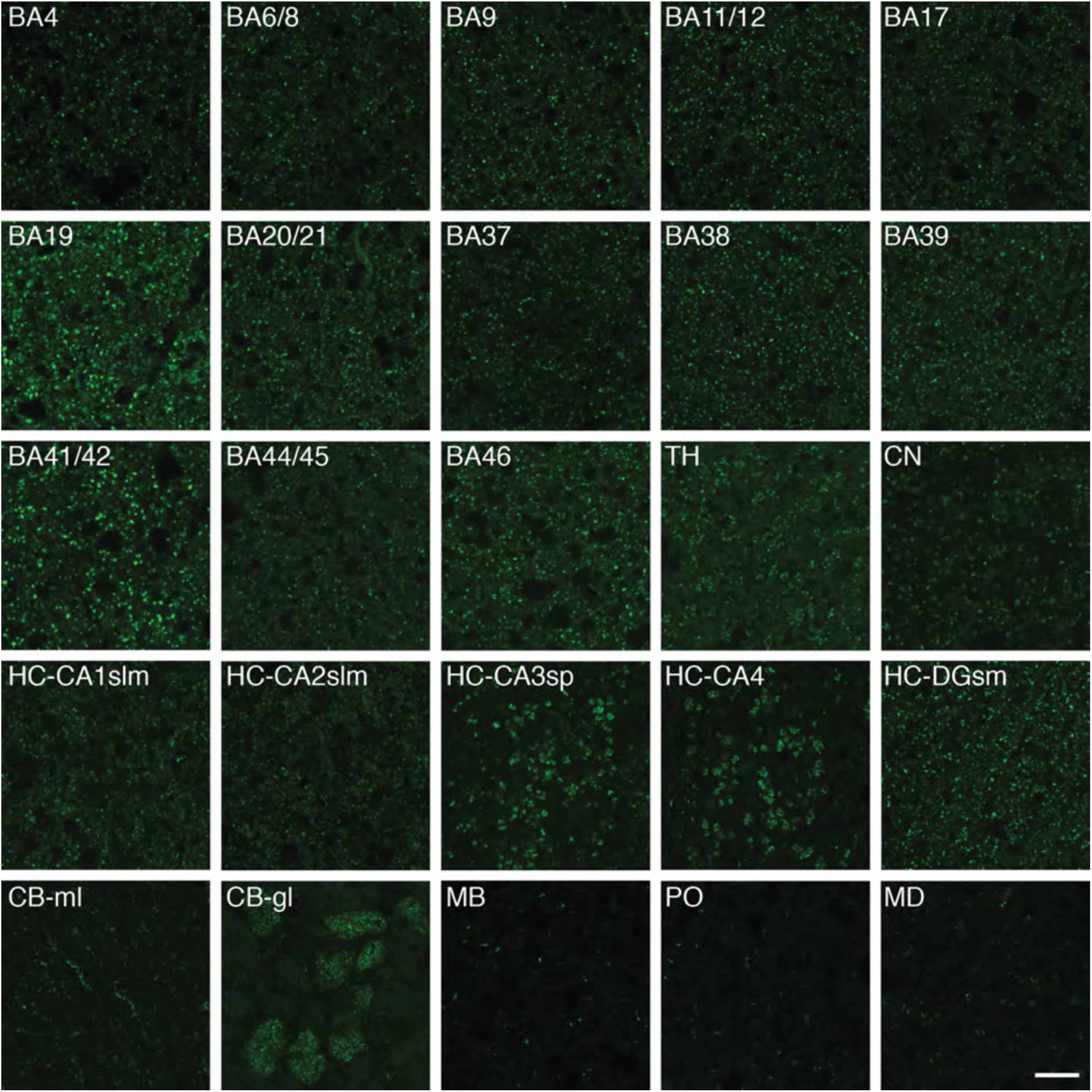
PSD95 expression in 20 human brain regions. Representative images from 20 human brain regions showing the different patterns of PSD95 distribution. BA, Brodmann area; TH, thalamus; CN, caudate nucleus; HC, hippocampus; CA, cornu ammonis; slm, stratum lacunosum – moleculare; sp, stratum pyramidale; DG sm, dentate gyrus stratum moleculare; CB, cerebellum; gl, granular cell layer; ml, molecular cell layer; MB, midbrain; PO, pons; MD, medulla. Scale bar: 10 μm.

In the human hippocampal formation, there was intense PSD95 staining within CA1-4 subregions as well as the dentate gyrus (DG) (Figure 6A-C). The highest intensity of staining was seen in CA1 (Figure 6D-I). Bright PSD95 staining was mostly observed within the dendritic layers, including stratum radiatum and stratum lacunosum-moleculare (Figure 6J) as well as the molecular layer of the DG (Figure 6K, S). Staining was observed on the cell bodies of the pyramidal cells of the CA1-4 subregions (Figure 6L, M), in contrast to the granule cells of the dentate gyrus that lacked staining (Figure 6N, O). CA1 contained small bright PSD95 puncta (Figure 6P), whereas CA2 puncta were dimmer (Figure 6Q). The puncta observed in the neuropil of the pyramidal cell layer of the CA3 (not shown) and CA4 subregions (Figure 6R) appeared very similar in size, intensity and pattern of staining: they were large, moderately bright, and formed a rosette-like pattern of staining.

**FIGURE 6.**
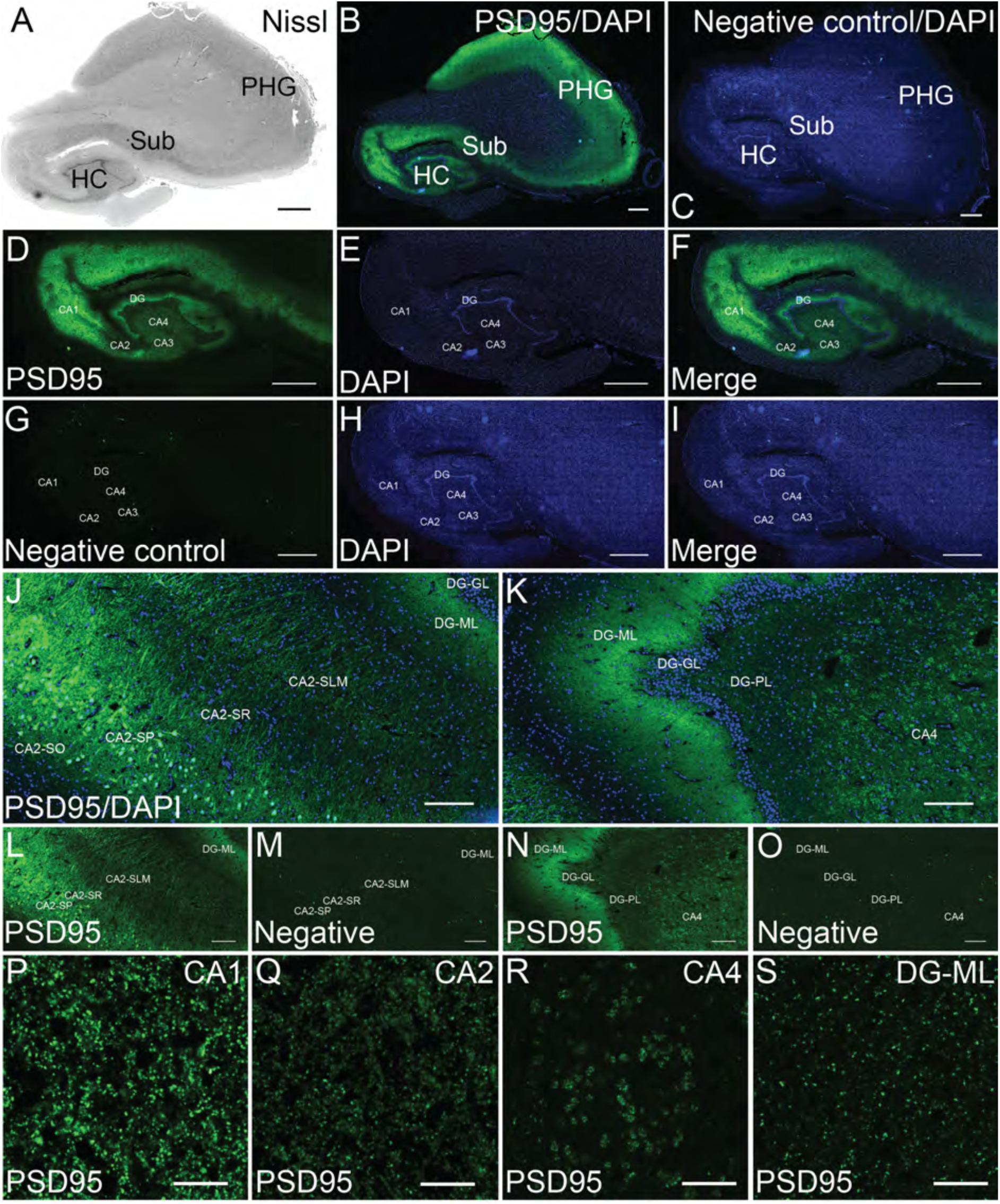
Pattern of PSD95 staining in the human hippocampus. (A) Nissl-stained section assessing hippocampal neuroanatomy. (B) PSD95 labelling in the hippocampus, subiculum and adjacent parahippocampal gyrus. (C) Negative control stained without primary antibody showing absence of PSD95 signal. (D) PSD95 labelling, (E) DAPI, and (F) merged imaged in the hippocampus proper. (G) Negative control, (H) DAPI, and (I) merged imaged in the hippocampus proper. (J) PSD95 labelling in the CA2 subregion. (K) PSD95 labelling in the DG subregion. (L) PSD95 labelling with DAPI, and negative control (M) in the CA2 subregion. (N) PSD95 labelling with DAPI, and negative control (O) in the DG subregion. (P-S) PSD95 staining in various hippocampal subregions. HC, hippocampus; Sub, subiculum; PHG, parahippocampal gyrus; DG, dentate gyrus; DG-ML, DG-molecular layer, DG-PL, DG-pleomorphic layer; DG-GL, DG granule cell layer; SR, stratum radiatum; SO, stratum oriens; SLM, stratum lacunosum-moleculare. Scale bars: (A-I) 2 mm, (J-O) 200 μm, (P-T) 10 μm.

The caudate nucleus showed strong PSD95 labelling (Figure 7A-C), with small patches (Figure 7B asterisk) that we confirmed as striosomes by IHC with Calbindin-D28K (Ito *et al*., 1992) (Figure 7D-F), which are embedded within the matrix. In addition to dendritic staining, somatic staining was observed, presumably in medium spiny neurons (MSNs). In the thalamus, moderate numbers of PSD95 positive puncta were detected (Figure 7G-I). PSD95 puncta were often found lined up along neuronal dendrites (Figure 7J), although a more diffuse distribution of puncta was also seen.

**FIGURE 7.**
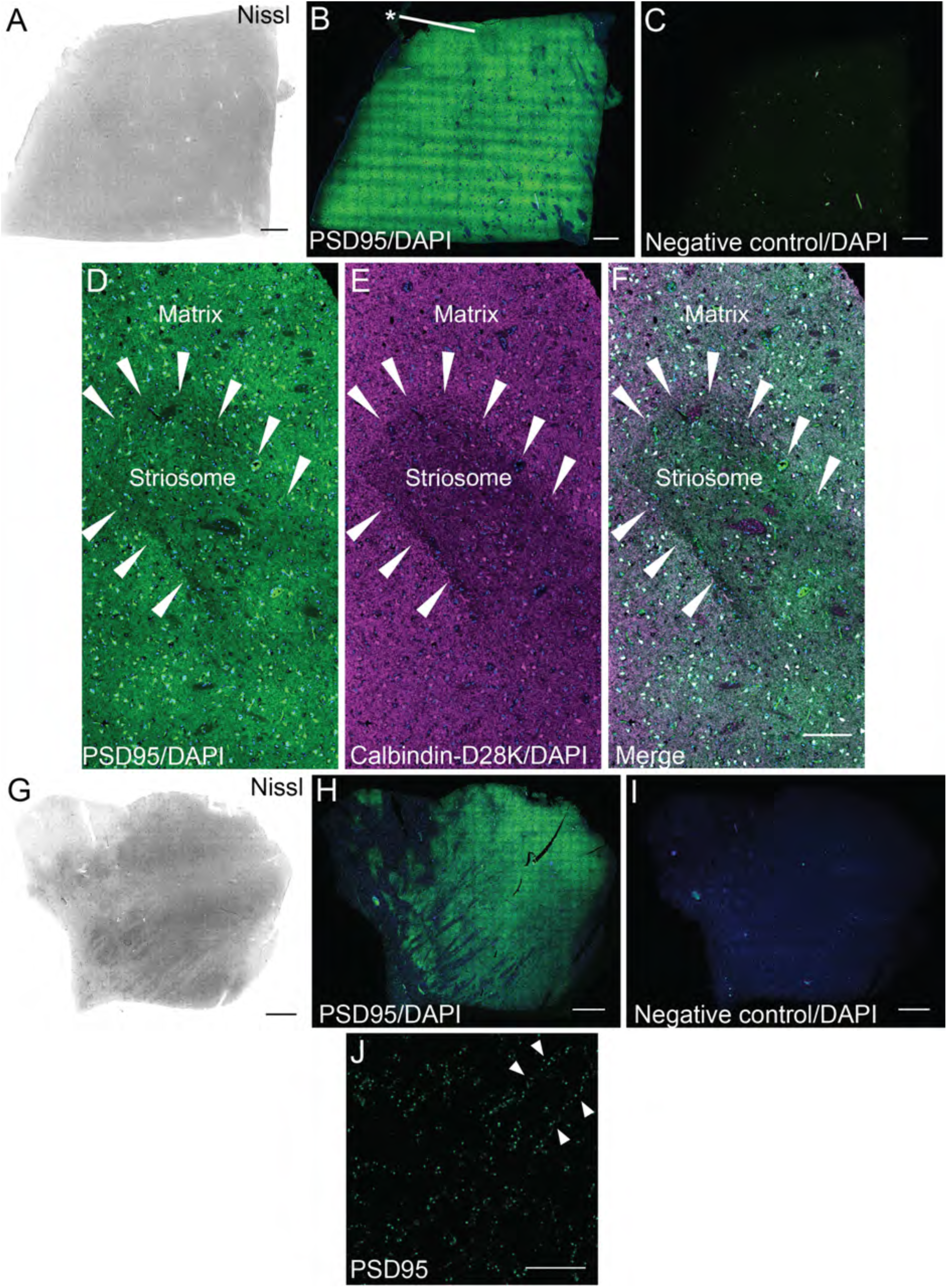
PSD95 staining in the human caudate nucleus and thalamus. (A) Nissl-stained section to assess neuroanatomy of the caudate nucleus. (B) PSD95 staining was not homogenous as small darker patches of striosomes were identified scattered within the caudate matrix (indicated with asterisk). (C) Omission of PSD95 primary antibody showed a lack of PSD95 labelling. (D) PSD95 puncta within a striosome (arrowheads) and a matrix. Somatic staining of PSD95 in medium spiny neurons was present in addition to dendritic PSD95 puncta. (E) The same striosome visualized with Calbindin-D28K antibody. (F) Merged image showing PSD95 and Calbindin-D28K labelling. (G) Nissl-stained section to assess thalamic neuroanatomy. (H) PSD95 labelling was variable, but prominent throughout the thalamus. (I) Omission of PSD95 primary antibody showed lack of PSD95 expression. (J) Punctate PSD95 staining along dendrites (arrowheads) within the thalamus. Scale bars: (A-C, G-I) 2 mm, (D-F) 100 μm, (J) 10 μm.

In the cerebellum PSD95 staining was restricted to the cerebellar folia, absent from the white matter and present at low levels in the dentate nucleus (Figure 8A-G). The adult cerebellar cortex is a three-layer structure composed of an internal granular cell layer (GCL), Purkinje cell layer (PCL) organized into a single row of Purkinje cells (PCs), and a molecular cell layer (MCL), and the pattern of PSD95 distribution varied between these layers (Figure 8H, I). Overall, PSD95 staining was higher in the GCL than in the MCL, and no PSD95 staining was seen in the PCs (Figure 8H, I). Numerous and dense clusters of PSD95 puncta were dispersed throughout the glomerular region of the GCL between the granule cells (Figure 8H small arrowheads). PSD95 puncta were only detected in the terminal pinceau (also known as the presynaptic plexus) of cerebellar basket cells, surrounding the axon hillock regions of PCs, but the postsynaptic PCs neurons did not express the protein (Figure 8H long arrowheads). A much lower density of PSD95 was detected within the neuropil of the MCL, where staining within the larger nuclei of the stellate and basket cells was observed (Figure 8H asterisks). Higher magnification images highlighted the clusters of PSD95 puncta within the GCL between the granule cells, corresponding to mossy and climbing fibers, but the granular cells only showed non-specific staining (Figure 8J). A much lower density of PSD95 was detected within the neuropil of the MCL, where PSD95 puncta formed a fine linear pattern along dendrites (Figure 8K). PSD95 puncta were not present in the white matter (Figure 8L).

**FIGURE 8.**
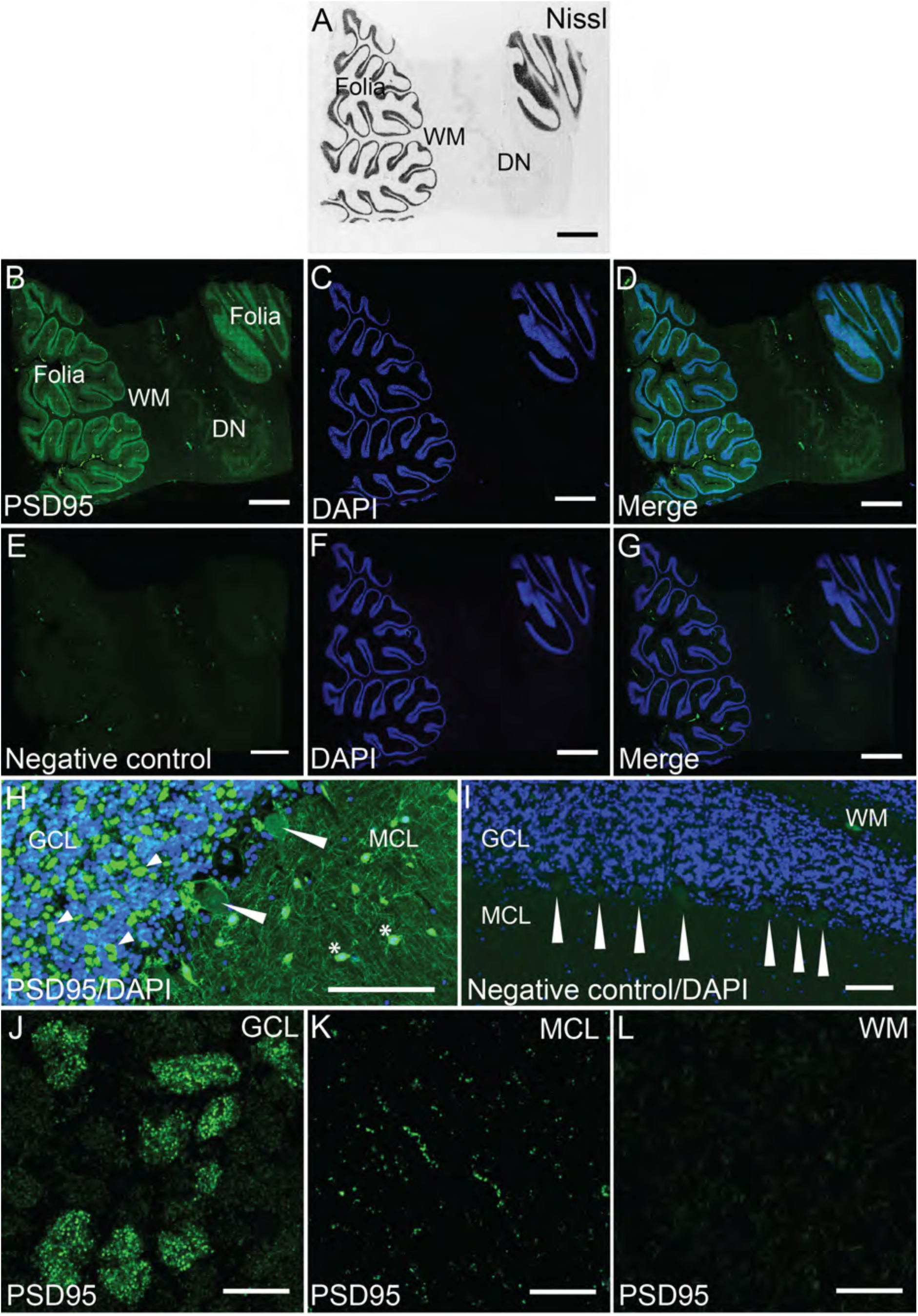
Pattern of PSD95 staining in the human cerebellum. (A) Nissl-stained section assessing cerebellar neuroanatomy: cerebellar cortex (folia), white matter and the DN. (B) PSD95 labelling in the cerebellar folia, white matter and dentate nucleus with (C) DAPI, and (D) merged images. (E) Negative control stained without primary antibody showing absence of PSD95 staining with (F) DAPI, and (G) merged images. (H) Numerous and dense clusters of PSD95 puncta dispersed throughout the glomerular region of the GCL between the granule cells (short arrowheads). PSD95 puncta were detected in the terminal pinceau of cerebellar basket cells, but PC neurons did not express PSD95 (long arrowheads). In MCL, PSD95 was observed as fine linear pattern with strong labelling of stellate and basket cell nuclei (asterisks). (I) PSD95 staining abolished in a negative control section, showing lack of staining around and within PCs (long arrowheads). (J) PSD95 staining in the GCL showing numerous intense puncta within the clusters. Arrowheads point to areas of rosette-like punctate formations. (K) PSD95 staining within MCL localized along dendrites. (L) No punctate PSD95 labelling was present in the cerebellar white matter. WM, white matter; DN, dentate nucleus; GCL, granular cell layer; MCL, molecular cell layer. Scale bars: (A-G) 2 mm, (H, I) 100 μm, (J-L) 10 μm.

The midbrain (Figure 9A-C), pons (Figure 10A-C) and medulla (Figure 11A-C) were examined for PSD95 expression, and overall showed a sparser density of fluorescent puncta than the supratentorial structures. Within the midbrain, a low density of PSD95 puncta was present within the substantia nigra (Figure 9D-F), and a lower density was detected in the periaqueductal grey area (Figure 9G-I). The fluorescent signals in these areas were moderately bright (Figure 9J-L). Within the pons, moderate density of PSD95 puncta was seen in the pontine nuclei (Figure 10D-F) and low density in the locus coeruleus (Figure 10G-I). The fluorescent signals in these pontine areas were moderately bright (Figure 10J, K). Finally, within the medulla, a moderate density of PSD95 puncta was present within inferior olivary nucleus (Figure 11D-F), and a low density was detected in the hypoglossal nucleus (Figure 11G-I). The PSD95 puncta in these medullary areas were dim (Figure 11J, K).

**FIGURE 9.**
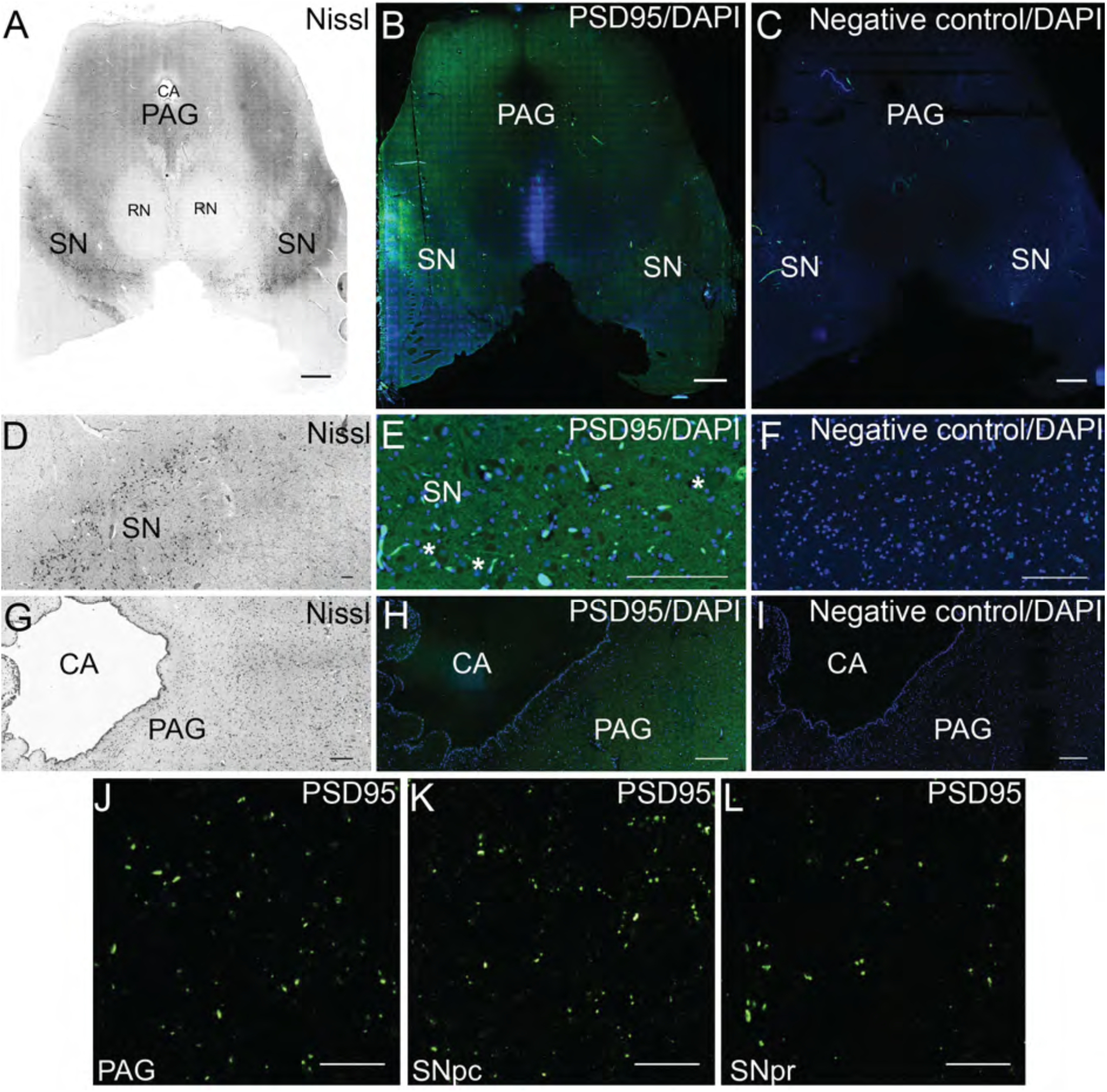
Pattern of PSD95 staining in the human midbrain. (A) Nissl-stained section to assess neuroanatomy of the midbrain. (B) Overview of PSD95 staining within the midbrain. (C) Omission of PSD95 primary antibody showed a lack of PSD95 labelling. (D) Nissl-stained section of the SN within the midbrain. (E) PSD95 staining within the SN showed low density of PSD95 expression. PSD95 labelling was absent from large pigmented neurons of the SN, as highlighted by asterisk. (F) No PSD95 expression was detected in the SN in the negative control. (G) Nissl-stained section of the PAG area within the midbrain. (H) PSD95 staining within the periaqueductal grey showed very low density of PSD95 expression. (I) Negative control showing no PSD95 expression within PAG. (J) Sparse density of PSD95 puncta in PAG. (K) Sparse density of PSD95 puncta in SNpc. (L) Sparse density of PSD95 puncta in SNpr. CA, cerebral aqueduct; PAG, periaqueductal grey; RN, red nucleus; SN, substantia nigra; SNpc, substantia nigra pars compacta; SNpr, substantia nigra pars reticularis. Scale bars: (A-C) 2 mm, (D-I) 100 μm, (J-L) 10 μm.

**FIGURE 10.**
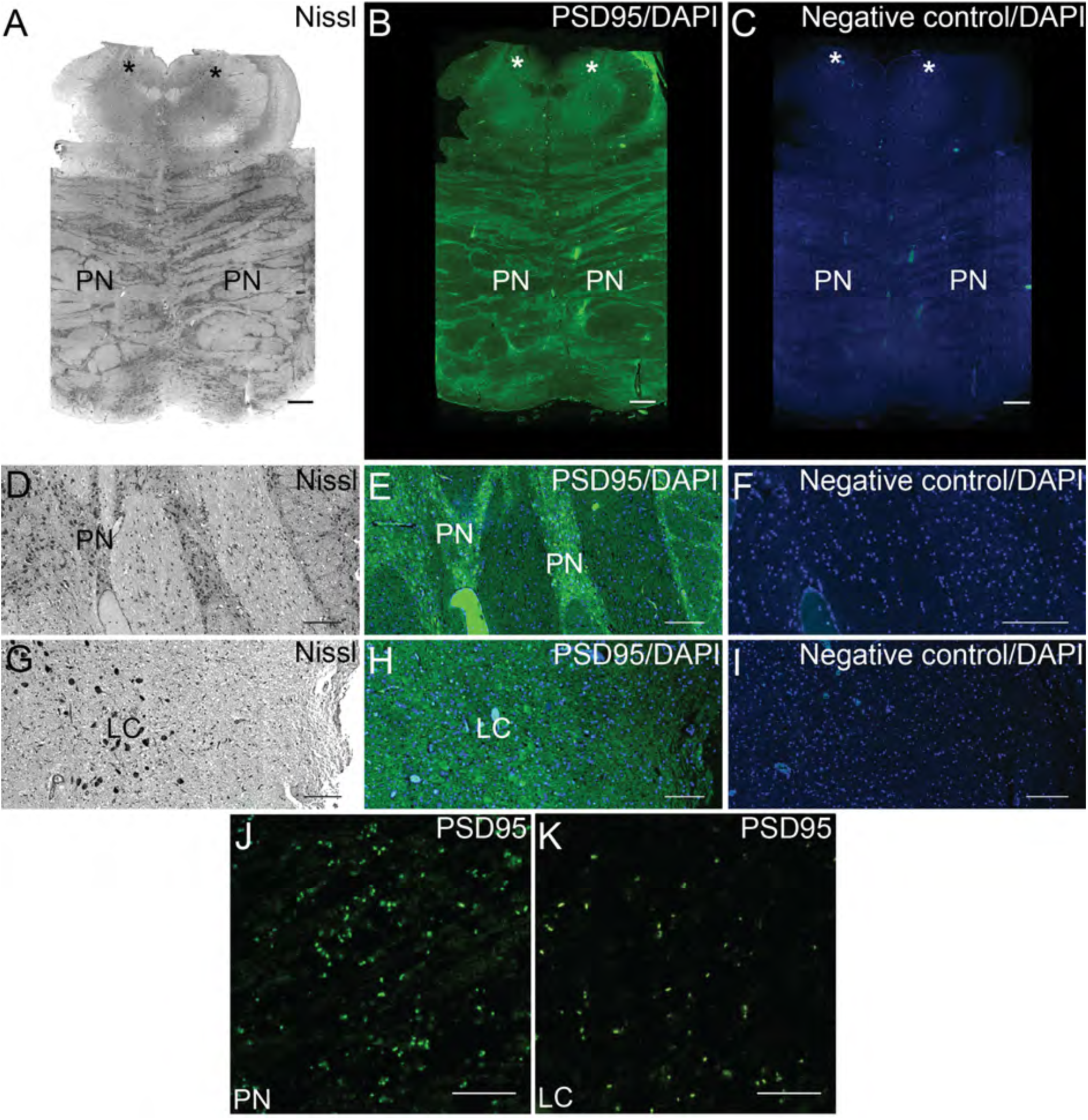
Pattern of PSD95 staining in the human pons. (A) Nissl-stained section used to assess neuroanatomy of the pons. LC is shown with asterisks. (B) Overview of PSD95 staining within the pons. (C) Omission of PSD95 primary antibody showed a lack of PSD95 labelling. (D) Nissl-stained section of the PN. (E) PSD95 staining within the PN showed low density of PSD95 expression. (F) No PSD95 expression was detected in the PN in the negative control. (G) Nissl-stained section of the LC within the pons. (H) PSD95 staining within the LC showed very low density of PSD95 expression. (I) Negative control showing no PSD95 expression within the LC. (J) Low density of PSD95 puncta in the PN. (K) Sparse density of PSD95 puncta in the LC. PN, pontine nuclei; LC, locus coeruleus. Scale bars: (A-C) 2 mm, (D-I) 100 μm, (J, K) 10 μm.

**FIGURE 11.**
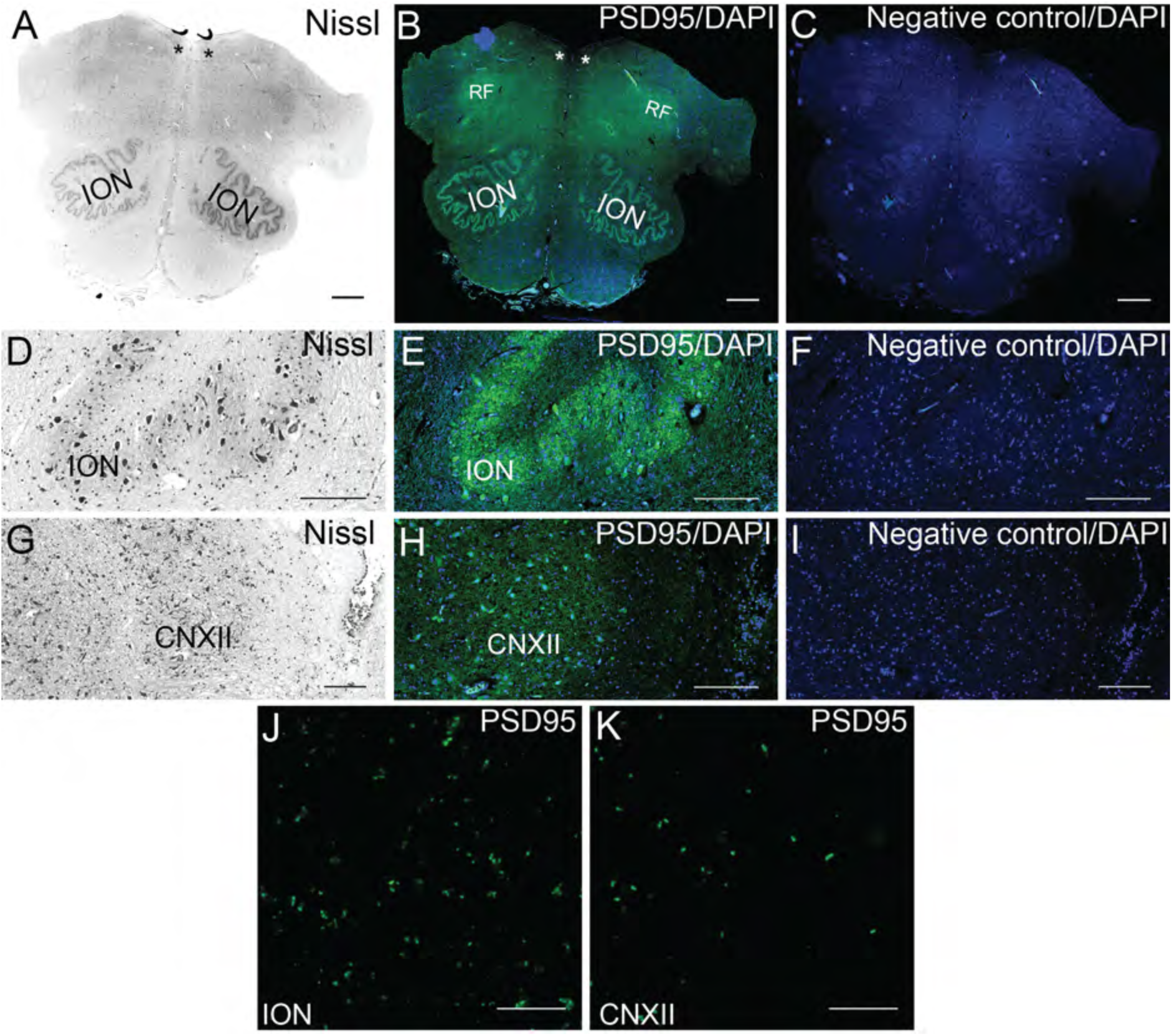
Pattern of PSD95 staining in the human medulla. (A) Nissl-stained section used to assess neuroanatomy of the medulla. Asterisks show location of CNXII. (B) Overview of PSD95 staining within the medulla. (C) Omission of PSD95 primary antibody showed a lack of PSD95 labelling. (D) Nissl-stained section of the inferior olivary nucleus. (E) PSD95 staining within the ION showed low density of PSD95 expression. (F) No PSD95 expression was detected in the ION in the negative control. (G) Nissl-stained section of the CNXII. (H) PSD95 staining within the CNXII showed very low density of PSD95 expression. (I) Negative control showing no PSD95 expression within the CNXII. (J) Low density of PSD95 puncta in the medullary ION. (K) Sparse density of PSD95 puncta in the CNXII. ION, inferior olivary nucleus; CNXII, nucleus of the cranial nerve XII (hypoglossal nucleus). RF, reticular formation. Scale bars: (A-C, G, H) 2 mm, (D-F), (I-K) 100 μm.

### Quantification of PSD95 puncta in human brain regions

To quantify synapses and describe their diversity in the brain regions we used the SYNMAP image analysis pipeline created to map the mouse synaptome (Zhu *et al*., 2018b). Each PSD95 punctum was quantified in terms of its intensity (a.u.) and size (μm^2^). In addition, the density of PSD95 puncta per 100 μm^2^ was calculated as a mean of the number of puncta per region. The mean values for four individuals are shown in Figure 12A-C and the data from individual subjects are shown in Figure S1. Figure 12D shows a summary of the mean values of each of the puncta parameters on maps of the human brain and highlights the diversity of these regions.

**FIGURE 12.**
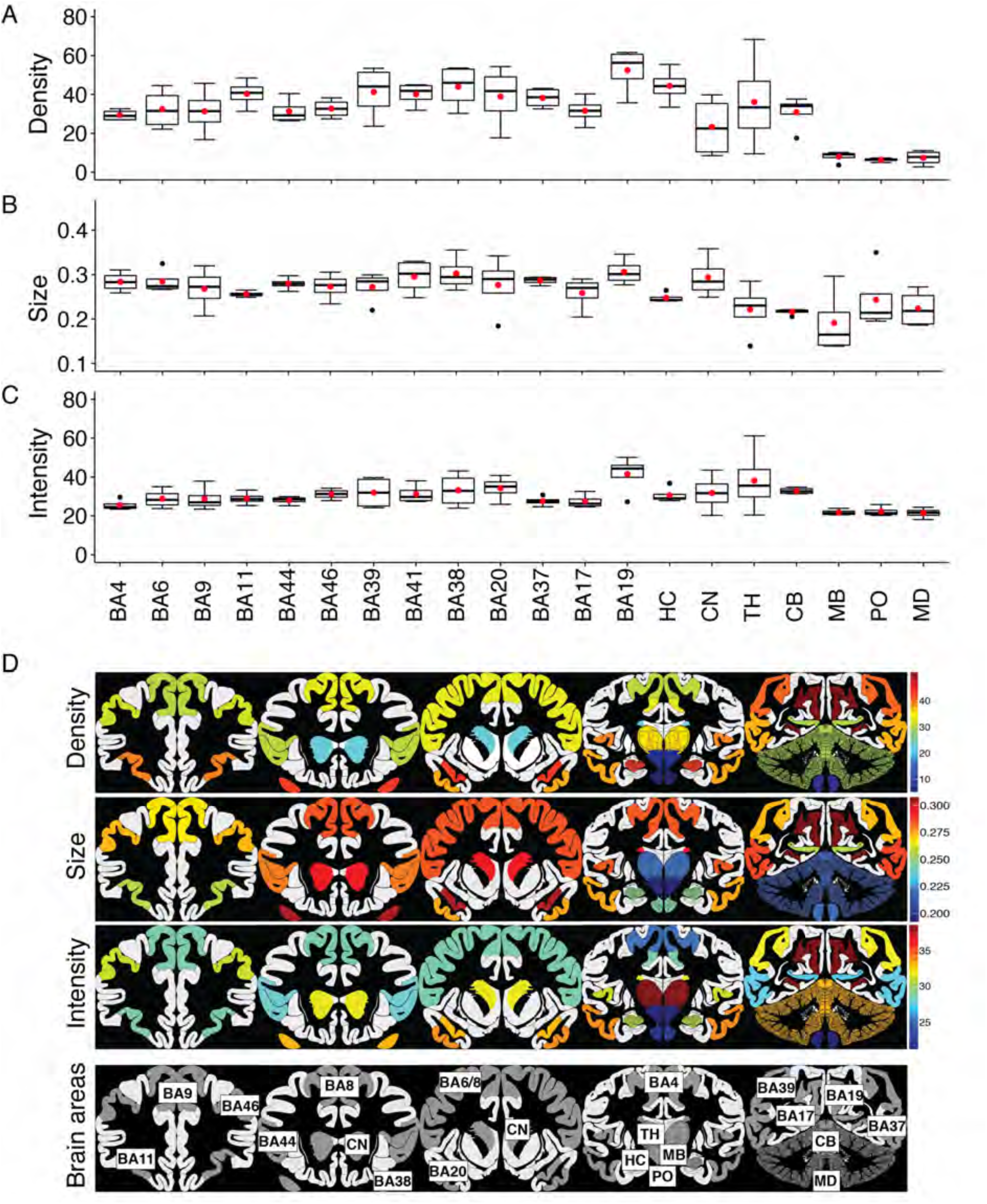
Quantification of PSD95 in 20 human brain areas. Boxplots showing quantification of PSD95 puncta in 20 human brain areas from four control cases. Data are presented as: (A) PSD95 punctum density, (B) PSD95 punctum size, and (C) PSD95 punctum intensity. (D) Colour-coded PSD95 synaptome map based on quantifications of parameters. Brain areas: BA, Brodmann area; HC, hippocampus; TH, thalamus; CN, caudate nucleus; CB, cerebellum; MB, midbrain, PO, pons; MD, medulla.

Ranking the 20 brain regions shows that the highest synapse density is in the neocortex and hippocampal formation and the lowest density in pons, midbrain and brainstem. There was a 7.3-fold difference between the highest and lowest mean density in the examined areas. Cortical area BA19, a region of the occipital cortex involved with feature extraction of visual images, had the highest density, intensity and size of synapses (Figure 12A).

Quantification of PSD95 puncta size across the 20 human brain areas is shown in Figure 12B. Size reflects PSD size within an individual synapse (Broadhead *et al*., 2016; Zhu *et al*., 2018a). There was a 0.5-fold change between the largest and smallest mean PSD95 puncta size. The secondary visual cortex, area BA19, had the largest PSD95 puncta, whereas midbrain had the smallest.

Ranking PSD95 puncta intensity across the 20 human brain areas is shown in Figure 12C. Intensity reflects the amount of PSD95 protein in an individual synapse (Broadhead *et al*., 2016; Zhu *et al*., 2018a). There was a 0.9-fold change between the highest and the lowest mean intensity in the examined areas. The secondary visual cortex, area BA19, showed the highest intensity of PSD95 puncta, whereas medulla showed the lowest. The deep grey nuclei (thalamus and caudate nucleus) showed greatest variations between individuals. Together these data show that PSD95 puncta parameters vary between the delineated 20 human brain regions (Figure 12D), with each brain area having a characteristic ‘signature’ of these parameters.

### The human synaptome architecture is hierarchically organized and similar to mouse

With the quantitative synaptic data in hand we could now make comparisons between human brain regions. We first examined the synaptome similarity of the human brain regions by generating a similarity matrix (Figure 13A), which revealed a number of interesting patterns. All neocortical areas and hippocampus showed high similarity (with the exception of BA19) and these regions were distinct to cerebellum and brainstem structures. BA19 is the visual association area and it has the highest PSD95 density, intensity and size. Interestingly, a very similar organization was observed in the mouse brain (Zhu *et al*., 2018a). To test if the synaptome architecture of the two species was conserved we correlated the synaptic puncta parameters between the homologous brain regions of the two species and found significant correlations (puncta density, r=0.90, p<0.0001; intensity, r=0.51, p=0.05; size, r=0.56, p=0.03)(Figure 13B).

**FIGURE 13.**
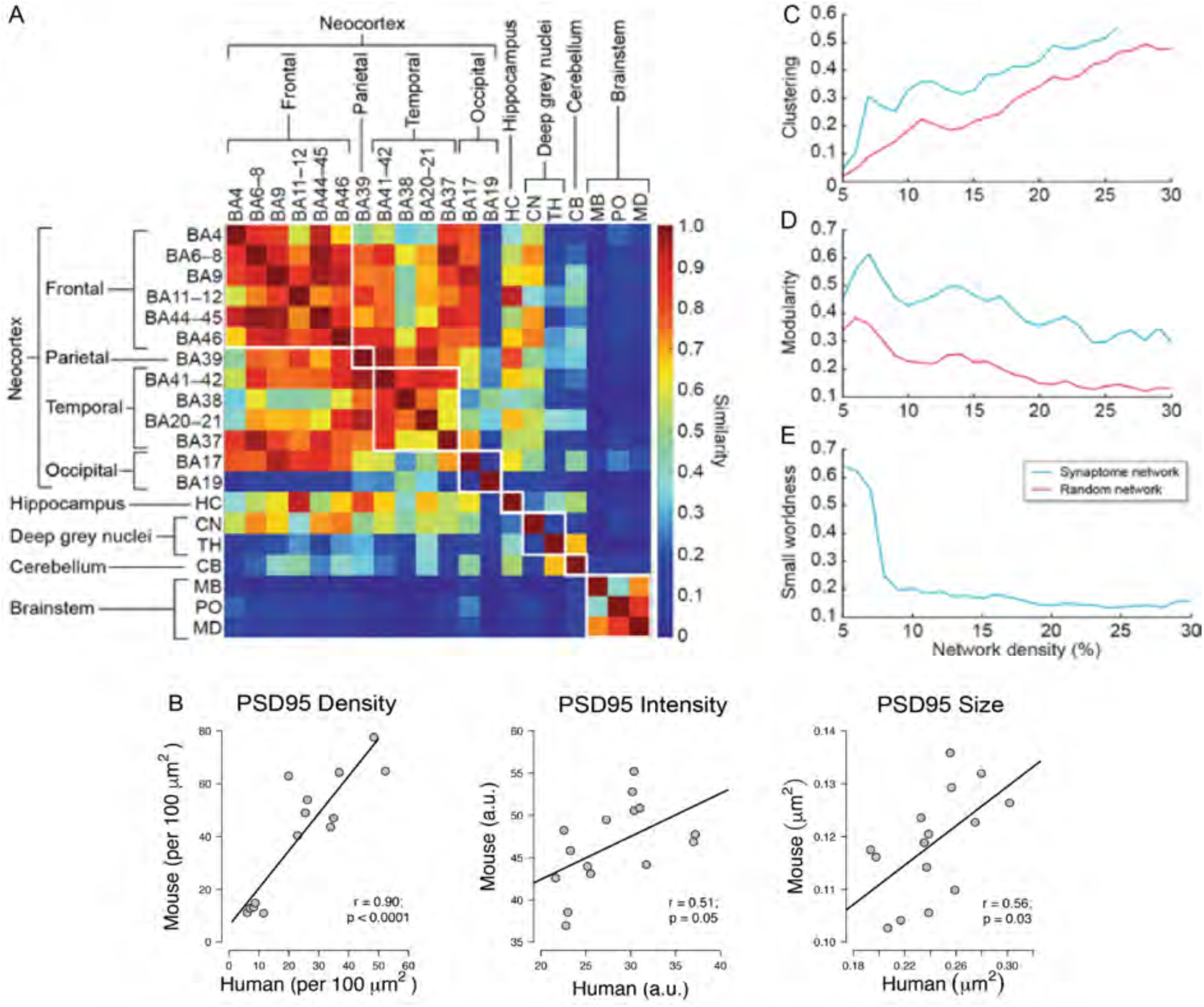
Synaptome architecture in human and mouse. (A) Similarity matrix between pairs of 20 regions (rows and columns). (B) Correlation of PSD95 positive puncta parameters quantification between human and mouse. Scatterplots showing correlations of PSD95 immunofluorescent puncta parameters (density, intensity and size) quantified from 15 brain areas of knock-in mouse and human tissue stained using PSD95 antibody. There was a high, positive linear and significant correlation between the human and mouse PSD95 puncta counts. A moderate positive, linear and significant correlation was found for PSD95 puncta intensity. PSD95 puncta sizes were moderately and significantly correlated between the species. Pearson’s product-moment correlation (r) and p values are provided. Values are based on three male WT mice aged 12 months and three male human controls. (C-E) PSD95 network topology. Clustering coefficient (C), modularity (D), and small-worldness (E) of the PSD95 network and random network.

The synaptome architecture can also be analyzed using network methods and in the mouse the synaptome shows a small-world network architecture (Zhu *et al*., 2018a). We analyzed the human synaptome network properties of the 20 human regions: each node in the network represented one region and edges that link nodes were scored for similarity of PSD95 parameters. Three network topological properties, clustering coefficient (CC), modularity and small-worldness (Figure 13C-E) were calculated and showed that the PSD95 network has a higher CC and modularity than the random network with equivalent network complexity. Thus, the PSD95 synaptome is considered to contain a small-world network structure with high small-worldness topology. Together these findings indicate that the human brain shares a synaptome architecture with similarity to mouse.

## Discussion

Using immunolabelling of PSD95 with high-resolution confocal microscopy together with the SYNMAP image analysis software, we have examined the distribution of excitatory synapses across 20 regions of the human brain in four individuals. We found that there are populations of PSD95+ synaptic puncta with varying size and intensity distributed in each brain region. Comparisons of the mean values of the synaptic parameters of these populations revealed a hierarchical organisation of the regions in which cortical and hippocampal regions shared greatest similarities, and cerebellar and brainstems had distinct synaptome signatures. This organisation is remarkably similar to the synaptome organisation in the mouse brain (Zhu *et al*., 2018a). Moreover, we found the synaptic puncta density, size and intensity correlated between homologous brain regions in mouse and human, showing evidence of a conserved synaptome architecture. Furthermore, the synaptome architecture of both species showed small world network properties. These data indicate that the synaptome architecture retained invariant features during the 90 million years since humans and mice shared a common ancestor and despite the fact that brain size is 1000-fold greater in humans. Previous studies have shown that the synapse proteome composition is also highly conserved between mouse and human (Bayés *et al*., 2011; Bayes *et al*., 2012) and regional diversity in the postsynaptic proteome of the human cortex (Roy *et al*., 2018a) and mouse brain shows similar features (Roy *et al*., 2018b).Thus, the molecular anatomy of synapse protein composition, synapse diversity and its spatial organization has conserved features between humans and mice.

The present study has been restricted to a single synaptic marker of excitatory synapses. Because high synapse diversity arises from the differential distribution of synaptic proteins and their cognate multiprotein complexes (Grant, 2018; Zhu *et al*., 2018a; Grant, 2019), it is likely that a detailed characterization of the synaptome of diverse molecular types of synapses in humans and mice will reveal species-specific synapses and synaptome maps. Understanding these differences could be extremely important for translational studies from the mouse. Mice carrying mutations in human orthologs that cause schizophrenia and autism have been shown to have altered synapse diversity and synaptome maps (Zhu *et al*., 2018a; Grant, 2019). The availability of synaptomic methods in humans now makes it possible to compare the synaptomes of the two species carrying the same mutations.

A limitation of this study is that we have not created a fine-scaled synaptome map of all the subregions within the 20 regions we have studied. This will require more extensive microscopy, scanning larger areas of brain tissue. We are currently designing experiments and developing technical approaches suitable for mapping all grey matter regions in whole coronal sections of human brain, which is a step toward a brain-wide synaptome atlas of the human. The availability of a human synaptome atlas for multiple synapse types and comparable atlases in other mammalian species will inform on the function of the human brain and the synaptomic basis of behaviour.

## Acknowledgements

OEC was supported by the Medical Research Council Clinical Research Training Fellowship - Scottish Clinical Pharmacology and Pathology Programme (SCP3). ZQ and SGNG are supported by the Wellcome Trust (202932) and the European Research Council (ERC) under the European Union’s Horizon 2020 research and innovation programme (695568 SYNNOVATE). The authors thank C.L. McLaughlin for technical support; J. Menendez Montes, J. Piatkowski for computational support; F. Zhu for mouse tissue samples; D. Maizels for artwork; C.S. Davey for editing. All human tissues were from the Edinburgh Brain Bank, funded by the Medical research Council (MR/L016400/1). We thank the patients and families who kindly donated the brains for this work and the staff at the EBB.

## Data accessibility

The authors confirm that all data underlying the finding are available and will be shared with the research community upon request.

## Author contributions

OEC - immunofluorescence, microscopy platforms, image capture, data analysis; ZQ - image puncta analysis (detection, segmentation, quantification) and network analysis; OEC, ZQ, CS, SGNG writing; CS, SGNG - direction.

## Abbreviations

BA: Brodmann area
CA: Cornu Ammonis
CA1-SML: CA1-Stratum Moleculare Lacunosum
CA1-SO: CA1-Stratum Oriens
CA1-SP: CA1-Stratum Pyramidale
CA1-SR: CA1-Stratum Radiatum
CA2-SML: CA2-Stratum Moleculare Lacunosum
CA2-SO: CA2-Stratum Oriens
CA2-SP: CA2-Stratum Pyramidale
CA2-SR: CA2-Stratum Radiatum
CA3-SL: CA3-Stratum Lucidum
CA3-SML: CA3-Stratum Moleculare Lacunosum
CA3-SO: CA3-Stratum Oriens
CA3-SP: CA3-Stratum Pyramidale
CA4: Cornu Ammonis 4
CB: Cerebellum
CN: Caudate Nucleus
CNXII: Hypoglossal Cranial Nerve
DG-PL: DG-Polymorphic Layer
DG-SG: DG-Stratum Granulosum
DG-SM: DG-Stratum Moleculare
DG: Dentate Gyrus
EBB: Edinburgh Brain Bank
FFPE: formalin-fixed paraffin-embedded
HC: Hippocampus
IHC: immunohistochemistry
ION: Inferior Olivary Nucleus
LC: Locus Coeruleus
MB: Midbrain
MD: Medulla
PN: Pontine nuclei
PO: Pons
PSD95: postsynaptic density protein 95
SN: Substantia Nigra
TH: Thalamus

---

**FIGURE S1.**
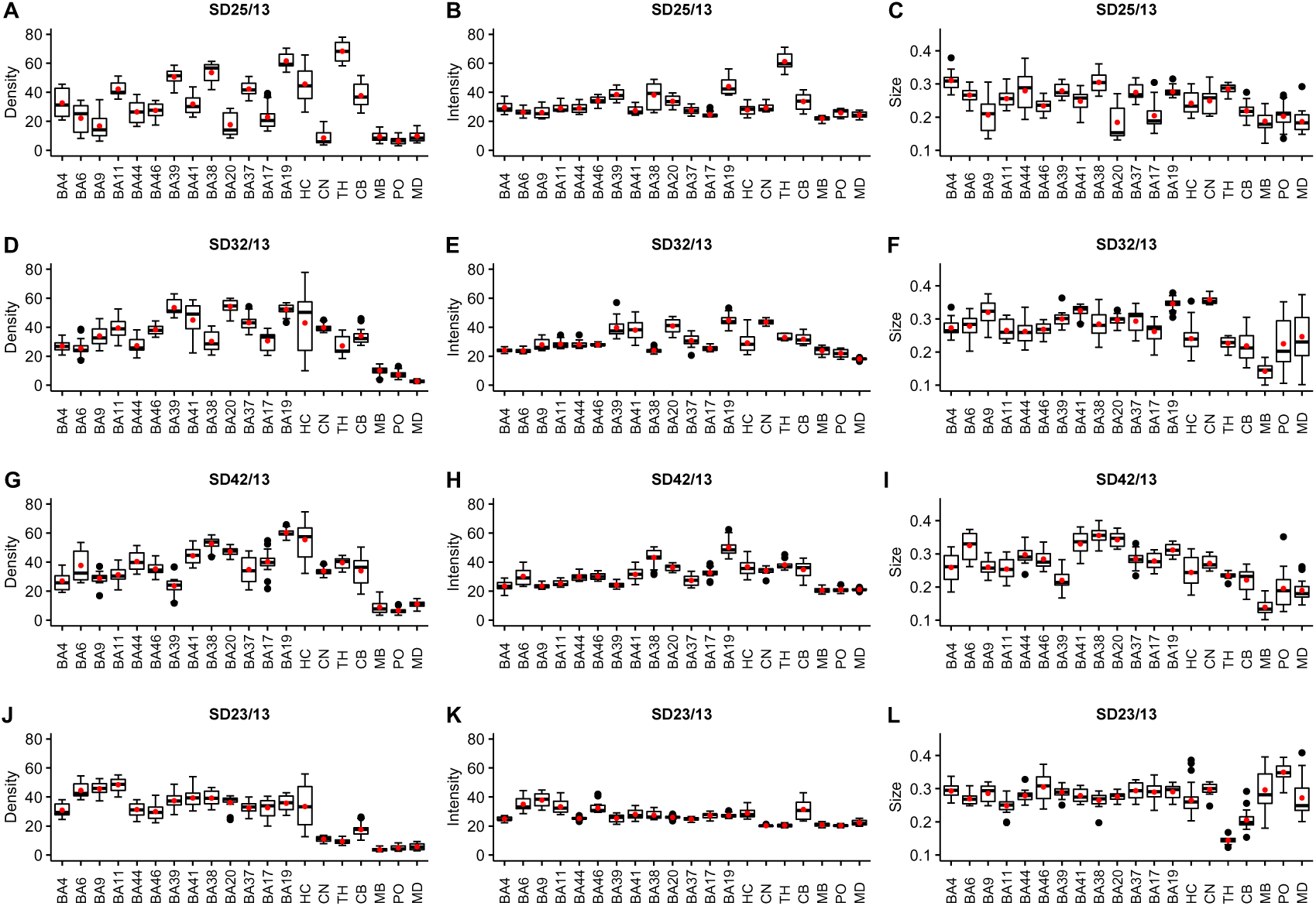
Comparison of PSD95 synaptic puncta parameter distribution from different brain areas for the four subjects studied. Cases: (A-C) SD25/13 (D-F) SD32/13 (G-I) SD42/13 (J-L) SD23/13 PSD95 puncta parameters: (A, D, G, J) PSD95 density (per 100 μm^2^) (B, E, H, K) PSD95 intensity (a.u.) (C, F, I, L) PSD95 size (μm^2^) BA, Brodmann area; HC, hippocampus; TH, thalamus; CN, caudate nucleus; CB, cerebellum; MB, midbrain; PO, pons; MD, medulla.

**TABLE S1.**
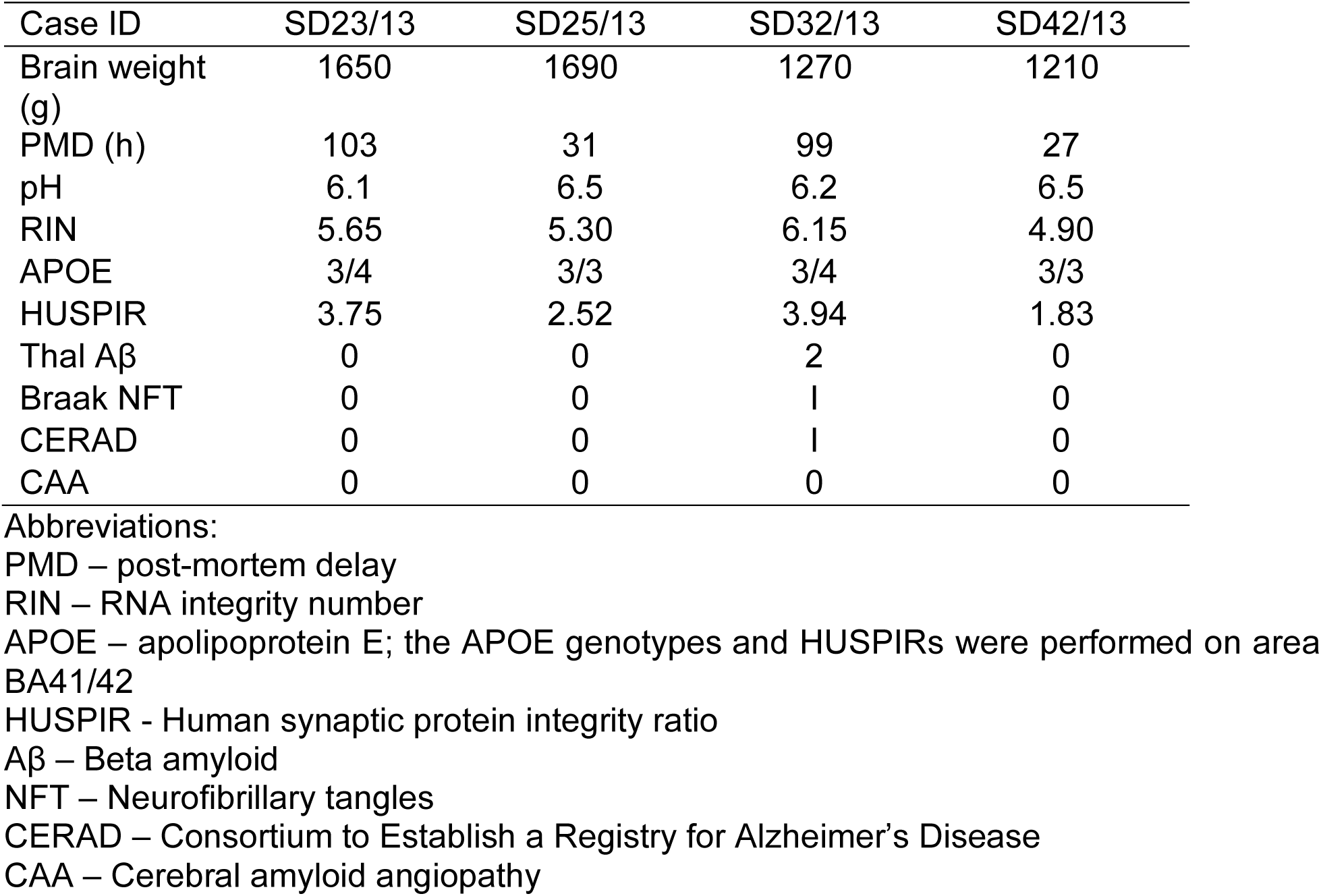
Neuropathological assessment of human tissue used in this study.

| Case ID | SD23/13 | SD25/13 | SD32/13 | SD42/13 |
| --- | --- | --- | --- | --- |
| Brain weight (g) | 1650 | 1690 | 1270 | 1210 |
| PMD (h) | 103 | 31 | 99 | 27 |
| pH | 6.1 | 6.5 | 6.2 | 6.5 |
| RIN | 5.65 | 5.30 | 6.15 | 4.90 |
| APOE | 3/4 | 3/3 | 3/4 | 3/3 |
| HUSPIR | 3.75 | 2.52 | 3.94 | 1.83 |
| Thal Aβ | 0 | 0 | 2 | 0 |
| Braak NFT | 0 | 0 | I | 0 |
| CERAD | 0 | 0 | I | 0 |
| CAA | 0 | 0 | 0 | 0 |

**Abbreviations:**
PMD – post-mortem delay
RIN – RNA integrity number
APOE – apolipoprotein E; the APOE genotypes and HUSPIRs were performed on area BA41/42
HUSPIR - Human synaptic protein integrity ratio
A $\beta$ – Beta amyloid
NFT – Neurofibrillary tangles
CERAD – Consortium to Establish a Registry for Alzheimer's Disease
CAA – Cerebral amyloid angiopathy

